# Structural flexibility of the human vault protein revealed by high-resolution cryo-EM and molecular dynamics simulations

**DOI:** 10.1101/2025.08.26.672097

**Authors:** Fabio Lapenta, Karen Palacio-Rodriguez, Sergio Cruz León, Simone Marrancone, Jana Aupič, Nils Marechal, Alexandre Durand, Dihia Moussaoui, Sonia Covaceuszach, Bhavani Gangupam, Claudia D’Ercole, Cristian Parra, Davide Cotugno, Giulia Tomaino, Paolo Tortora, Ario de Marco, Alberto Cassetta, Alessandra Magistrato, Gerhard Hummer

## Abstract

Vaults are massive ribonucleoprotein complexes, highly conserved and abundant in eukaryotic cells, yet with unclear function. Their thin-walled barrel-shape architecture is composed of two symmetrical, antiparallel half-shells, each containing 39 copies of the major vault protein (MVP). The spacious lumen of the vault suggests a role in cellular transport. To facilitate cargo encapsulation and release, the vault is thought to open into two halves, yet the molecular mechanism governing vault opening remains elusive. Here, we combine cryogenic electron microscopy (cryo-EM) and multi-scale molecular dynamics (MD) simulations to reveal the structural factors giving flexibility to the human vault protein. Using cryo-EM, we identified two alternative conformational states of the human vault, along with the half-vault shell. MD simulations of these structures show extensive, breathing-like motions, porous solvent-exposed surfaces, and distinct structural variability between conformational states. The stable intermediates and the flexibility at the interface of the half vaults together suggest a possible mechanism for the dynamic assembly and disassembly of the vault.

## INTRODUCTION

Vault particles are ubiquitous ribonucleoprotein (RNP) complexes with a distinct barrel-shaped architecture and a large lumen^1^. As the largest RNP yet described in eukaryotes, vaults are composed primarily of 78 copies of the 99 kDa major vault protein (MVP), self-assembled into a symmetric protein shell^2^. Pioneering structural studies on the vault from *Rattus norvegicus* revealed that two 39-mer halves of the vault particle establish an anti-parallel association in the central area of the barrel via their N-terminal domains. This assembly forms a particle with a large inner volume of around 40,000 nm^3^ and dimensions of 40 nm × 40 nm × 67 nm^3–6^ (**Figure 1A**). The unique architecture of the vault is inherently tied to the structure of MVP, with each MVP subunit consisting of 12 distinct domains (**Figure 1B**). Nine N-terminal antiparallel three-stranded β-sheet repeat domains (R1-R9) extend from the midsection and compose the central body of the vault^6^. The subsequent shoulder domain, with its globular α/β fold, is homologous to the Stomatin, Prohibitin, Flotillin, and HflK/C (SPFH) domain, characteristic of vault-like membrane-associated protein assemblies^7,8^ and forms a hinge between the central body and the C-terminal caps^9^. The latter are formed by long helical domains known as the cap-helices, which are engaged in an extensive coiled-coil helical bundle that stabilizes the vault^3^. Finally, at the two tips of the barrel, the C-terminal regions of MVP form the cap-ring, whose structure, resolved only at low resolution, appears partially disordered, possibly folding inwards^6,10^.

**Figure 1.**
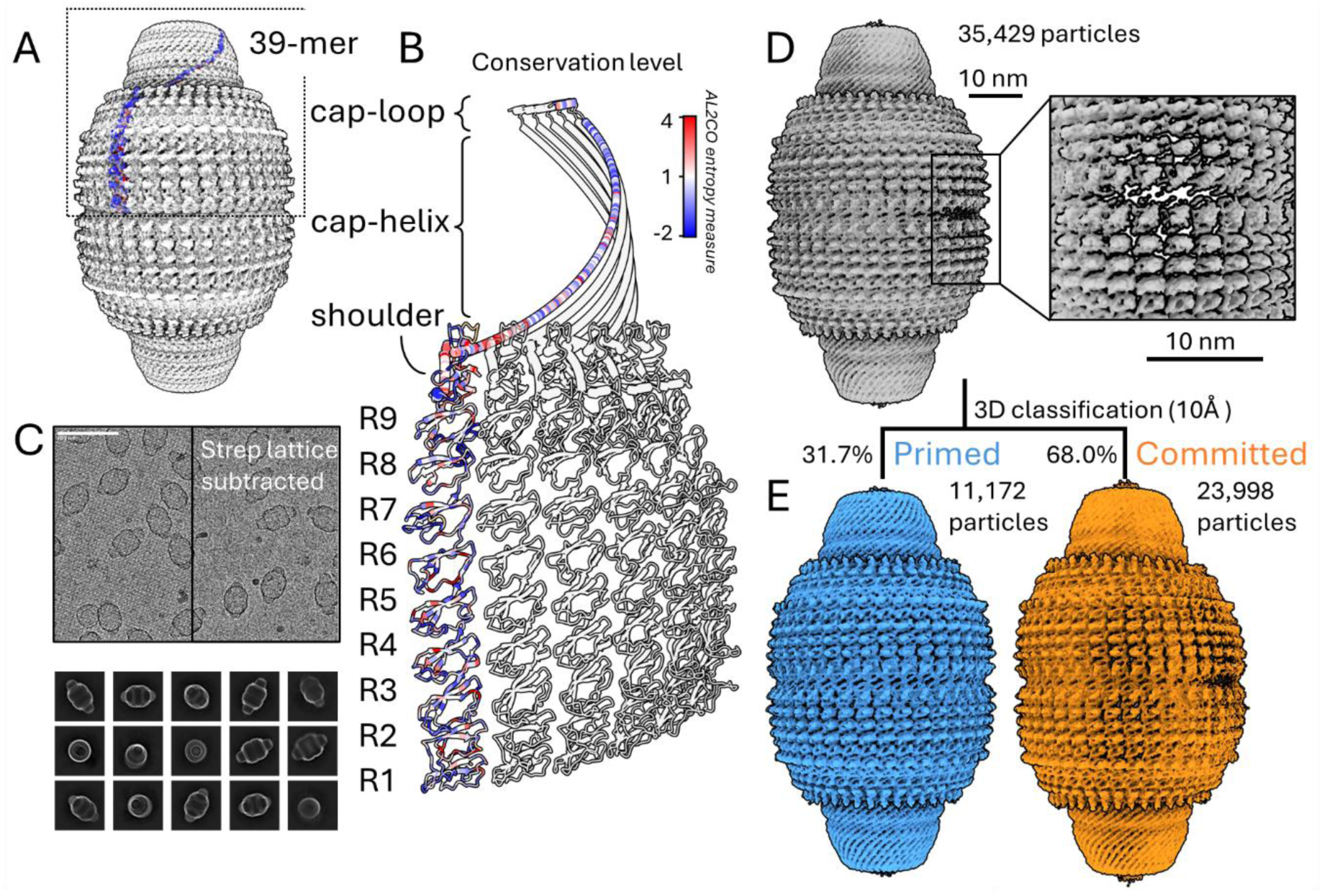
Major Vault Protein assembled into the 78-mer vault. A) Structure of the vault composed of 78 MVP copies. MVP monomer is coloured according to residue positional entropy-based conservation scores, calculated by AL2CO^44^. B) Detailed view of MVP monomers, highlighting individual domains: repeat domains (R1 to R9), shoulder domain, cap-helix, and cap-loop (PDB:4HL8)^10^. C) Top: micrograph before and after subtraction of the streptavidin 2D crystal lattice by Fourier filtering of Bragg spots. Bottom: 2D class averages of the vault selected for further refinement. D) Volume map of the initial density reconstruction obtained without imposing symmetry, based on 35,429 particles. The inset shows the symmetry-mismatched component of the map from panel D (60° rotated). E) Cryo-EM density maps obtained from 3D classification of the particles with filter resolution of 10 Å, which shows the overall structural differences between the two conformations.

Vaults are highly conserved RNPs with a nearly identical morphology across eukaryotes^11,12^. The high energetic cost of synthesizing vaults for the cell and their evolutionary conservation across species imply a pivotal role these RNPs play in the cellular environment. Nevertheless, while vaults have been implicated in numerous cellular processes (i.e. intracellular transport^13–15^ and signalling^16^, innate immune response^17,18^, DNA damage repair^19^, and apoptosis^20,21^, among others) their primary function remains unclear^22,23^. Vaults also have a long-debated role in multi-drug resistance^24–27^, neurodevelopmental disorders^28^ and recent findings have suggested their possible involvement in protein folding^29^. Macromolecular partners associated with vault particles *in vivo* include the vault-associate protein telomerase TEP1^30^, the poly-(ADP-ribose) polymerase vPARP^31^ and vault-specific short un-translated RNAs (vtRNA). Recently, electron-tomographic imaging of eukaryotic cells revealed vault particles capable of accommodating ribosomes within their inner cavity^32^, advocating for their ability to encapsulate also other, very large, cellular components. However, the purpose of compartmentalising molecular components within such a wide inner cavity is still obscure, compounding to the elusive cellular role of the vault. Interestingly, there is evidence that vPARP can enter pre-assembled vaults via a specific domain coined INT, suggesting that vaults are indeed dynamic assemblies capable of undergoing the conformational changes needed to house diverse molecular partners^31,33^. These examples point to a dynamic exchange of molecular partners between the vault’s interior and exterior. Accordingly, it was proposed that the vault may act as a shuttling scaffold for intracellular transport^33^. Indeed, it has been observed a dynamic exchange between the 39-mer halves composing the vault^34^. However, the molecular details of the opening and closing process and of its regulation in the cellular environment remain to be elucidated. Notably, it has been reported that vaults are sensitive to low pH, which weakens the interaction between MVP monomers^35^ and promotes vault dissociation into halves^36^.

X-ray structural characterization of the isolated N-terminal R1-R7 domains demonstrated that they adopt a more relaxed conformation – with a lower degree of curvature – compared to when integrated in the full-length, assembled vault^10,37^. Due to the large size of the particle itself, such conformational changes could be the result of intrinsic breathing motions that introduce flexibility and structural variability within the assembly, potentially leading to dissociation of the two halves. Indeed, 3D classification of cryogenic electron microscopy (cryo-EM) maps of the vault from *R. norvegicus* could distinguish regularly assembled, symmetric, vault particles from quasi-symmetric vaults distorted at the waist, thus capturing a partially open state of the vault^38^. This suggested that the vault opening is a multi-step process that occurs through a series of intermediate states, initiated by a prominent structural alteration at the N-terminal waist region. However, the low resolution of the reconstruction prevented a more detailed mechanistic analysis of the disassembly process.

In this work, we investigated the structure and dynamics of the human vault by cryo-EM and molecular dynamics (MD) simulations. Cryo-EM analysis captured vaults in two distinct conformations, one symmetric and the other asymmetric and apparently committed to opening. Atomic-level structures were determined and used as starting points for MD simulations. These simulations confirmed the stability of the two conformations, revealed mechanisms for passive diffusion of small molecules in and out of the vaults, and shed light on how vault dissociation is triggered in molecular detail. Together, our results uncover what appear to be the initial steps of human vault disassembly at atomic resolution.

## RESULTS

### MVP purification and cryo-EM imaging

Vault particles were expressed and purified from the yeast *Komagataella phaffii* (also *Pichia pastoris*). This organism lacks an endogenous vault gene in its genome and is characterized by rapid growth and stable gene expression^39^. We relied on a strain of *K. phaffii* bearing an integrated human MVP gene (hMVP) under the control of a constitutive promoter (pGAP), allowing intracellular expression of hMVP. The purification procedure, based on previously reported works^40,41^, consisted of an ultra-centrifugation step followed by RNase treatment prior to size-exclusion chromatography (SEC). The protein was analyzed by multi-angle-light scattering coupled to size exclusion chromatography (SEC-MALS) to assess the correct assembly of the complex (**Supplementary Figure 1**). The molecular weight of the main peak (7.66 ± 0.6% MDa) confirmed correct self-assembly of the particle in solution. However, batch small angle X-ray scattering (SAXS) analysis revealed the presence of both 78-mer vault and 39-mer half vaults in the sample (56% and 44%, respectively), consistent with the equilibrium between association and dissociation of half vaults reported in previous works^5,34^ (**Supplementary Figure 2A**).

The protein was biotinylated and subsequently immobilized on streptavidin-coated affinity grids to increase local concentration and overcome issues with preferred orientation of the particles on the grid, then vitrified and imaged using a 300 kV cryogenic electron microscope (**Supplementary Table 1**). Although the majority of the vault appeared intact on the grid (**Figure 1C**), micrograph analysis showed a heterogeneous population of vault particles in different states: whole vaults, half-vaults, and collapsed particles. For single particle analysis (SPA), we considered only intact vault particles, whereas half-vault and collapsed vault particles were identified and excluded (**Supplementary Figure 3**). A further selection was performed based on 2D classes averaging (**Figure 1C**).

Initial 3D refinement without enforced symmetry revealed a density map exhibiting the characteristic barrel-shaped vault architecture with a 39-fold dihedral quasi-symmetry, albeit with a visible distortion detected at the midsection of the vault (**Figure 1D**). The lower density observed in the main symmetry-mismatched component of the map suggested a degree of flexibility at the vault’s waist (**Supplementary Figure 4**). To evaluate the structural flexibility of these particles, we performed three-dimensional variability analysis (3DVA) across the whole particle stack^42^, including particles from both conformations. 3DVA identified significant variability, revealing continuous flexibility within the vault, characterized by different modes of stretching and compression across the whole particle, which resulted in the formation of multiple symmetry-mismatch components in the committed conformation, with the most prominent distortion found at the waist (**Supplementary Movie 1** and **Supplementary Figure 4**).

To further investigate this heterogeneity, we performed an unsupervised 3D classification of the particles^43^ (filter resolution 10 Å), identifying two major sub-populations, distinguished by presence or absence of the D39 point-group symmetry that is a characteristic feature of the vault particle. In line with a similarly processed cryo-EM dataset of the vault from *R. norvegicus*^38^, where a distorted conformation of the particle was detected, we named these subpopulations as *primed* (symmetric) and *committed* (asymmetric) conformations (**Figure 1E**). While these two conformations were distinguishable from the cryo-EM dataset, SAXS analysis was not able to discriminate between them due to their similarity in terms of overall size and molecular shape (**Supplementary Figure 2B**).

### Structural description of the primed vault

Single particle reconstruction of the primed conformation was carried out imposing 39-fold dihedral symmetry, resulting in a high-resolution refinement of the vault at 3.09 Å resolution, as determined by gold-standard Fourier shell correlation (GS-FSC) at an FSC threshold of 0.143 (**Supplementary Figure 5**). The lower density region at the N-terminal waist was improved by local refinement, yielding an additional map with GS-FSC resolution of 3.53 Å. This refinement provided better density for constructing the R1-R2 repeats in the atomic model (**Supplementary Figure 5**). This approach enabled tracing of the carbon backbone from residue 3 to residue 814, with the notable exception of two flexible loops (residues 429-449 and 608-619), which were also missing in previously published structures of MVP from *R. norvegicus* (rMVP). The map showed clear density signals for the sidechains along the whole sequence of the human MVP (hMVP) (**Figures 2A and 2B** and **Supplementary Figure 6**).

**Figure 2.**
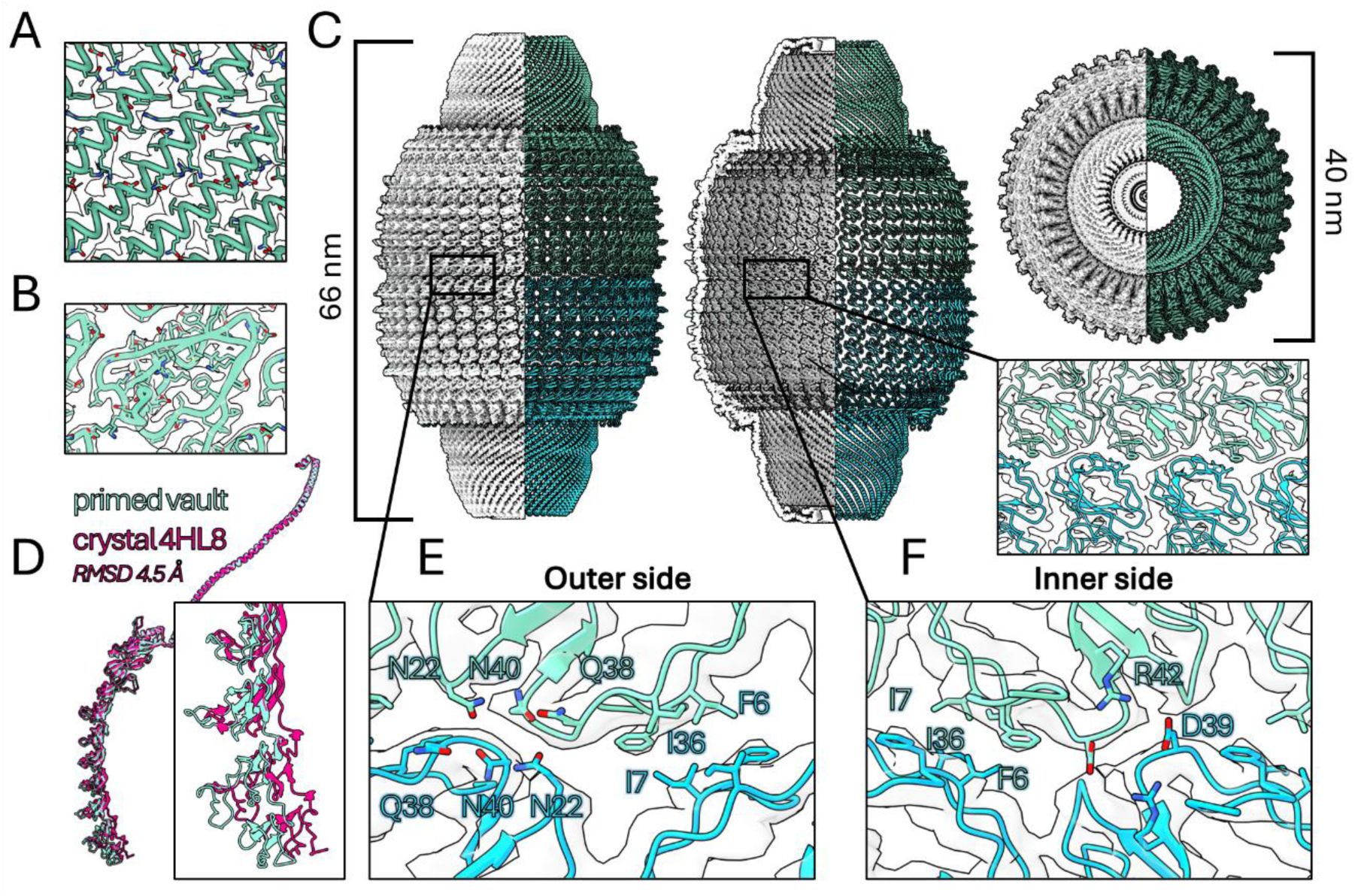
**Structural details of the vault structure in primed conformation**. A, B) Close-up views of the fit between the atomic model and the map for the cap-helix (A) and the R5 domain (B). C) Vault structure in primed conformation. The map is cropped in half longitudinally, with the corresponding atomic model showed side-by-side. The left panel shows the intact vault. The model in the central panel is clipped in the middle. The right panel shows the top view of the protein. The inset shows a detail of the fit between the map and the atomic model, represented as cartoon, at the midsection where the two 39-mer halves meet. D) Structure alignment of hMVP (from this work) and rMVP structure obtained by X-ray crystallography (PBD: 4HL8)^10^, in green and pink, respectively. The detail in the panel shows the R1-R4 domains. E) Close-up of the outer side of the vault waist in the primed conformation (local refinement map), showing the polar triad (Asn22, Gln38 and Asn40) and the hydrophobic patch (Phe6, Ile7 and Ile36). F) Close-up of the inner side of the vault waist in the hMVP structure (local refinement map). The salt bridge between Asp39 and Arg42 and the hydrophobic patch (Phe6, Ile7 and Ile36). All density maps are displayed at contour level of 0.8.

The reconstructed atomic model presented the characteristic vault architecture, with a shell composed of 78 repeated structural units of MVP, tightly interacting around a large inner cavity (**Figure 2C**). As expected from the high sequence homology (91.06%) between rMVP and hMVP, the structure of hMVP bore high similarity to previously deposited structures of its murine counterpart. The root mean square deviation (RMSD) between primed hMVP and primed rMVP (PDB: 7PKR)^38^, also resolved by cryo-EM, corresponded to 1.07 Å (**Supplementary Figure 7**). Conversely, RMSD (4.47 Å) was significantly higher when comparing to rMVP resolved by X-ray crystallography (PDB: 4HL8)^10^ (**Figure 2D**). The largest structural divergence between hMVP and crystallized rMVP was observed in the curvature of the N-terminal R1-R5 domains, a difference likely due to the absence of crystal packing contacts, as suggested in a previous work^9^.

A detailed analysis of the human vault in the primed conformation enabled the identification of interfacial contacts crucial for the assembly of MVP chains into the vault particle. The N-terminal R1 domains converge at the interface of the two 39-mer halves, establishing a network of interactions that tie the antiparallel halves together at the midsection of the vault. Each R1 domain interacts with two opposing R1 domains, forming an interface characterized by a hydrophobic patch composed of residues Phe6, Ile7, and Ile36 from two interacting chains. These residues form a tightly packed hydrophobic cluster, surrounded by polar amino acids that further stabilize the interaction between the two antiparallel R1 (**Figure 2E**). Additionally, a network of polar interactions is established along the outer side of the vault, involving amidic residues (Asn22, Gln38 and Asn40) from opposing R1 chains (**Figure 2E**). Likewise, inter-chain salt bridges between Asp39 and Arg42 of opposing chains propagated along the inner waist of the vault (**Figure 2F**). The lateral contacts between MVP monomers primarily reside in the helix-cap region, with 132 residues involved in neighbouring contacts between repeat domains and shoulder domain (residues 1-646), compared to 144 interacting residues in the helix-cap (residues 647-814) (**Supplementary Figure 8** and **Supplementary Figure 9**). Analysis of lateral contacts within the repeat domains identified Arg9 involved in hydrogen bonding with Gln21 of the neighbouring chain, along with several ionizable residues and six interfacial histidine residues (His85, His279, His356, His464, His534, and His592) likely involved in a hydrogen bonding network (**Supplementary Figure 8**).

The helix-cap stabilizes the half-vaults by establishing a wide 39-mer helix bundle. Analysis of the residue composition at the helix interfaces revealed an abundance of alanine and leucine residues densely packing the helices close together, in addition to charged residues on the outer position of the helix ridge (**Supplementary Figure 9**).

### Structural description of the committed vault

Unsupervised 3D classification of the particles identified a population of intact vaults in the micrographs, maintaining a discernible overall barrel-like architecture, albeit not fully symmetric. We designated this conformation as *committed* (**Figure 1E**). Refinement of the volume (without imposing symmetry) yielded a map with a dominant D39-point symmetry, similar to the primed conformation, yet with a clear symmetry-mismatched component, most prominent at the waist. The quasi-symmetrical nature of this assembly implied a high degree of structural flexibility in this conformation of the vault. Therefore, we performed a further refinement employing symmetry relaxation, a method that only limits the pose search for each particle to the original symmetry group of the assembly for the purpose of an asymmetric refinement^45^. This approach yielded a map at 4.45 Å GS-FSC resolution (**Supplementary Figure 10A**).

The map showed the vault committed conformation being overall comparable to the primed counterpart, with major differences clearly noticeable at the waist, corresponding to the N-terminal R1-R6 domains of 8 chains in each of the two 39-mer halves (**Figures 3A** and **3B**). This deformation was more pronounced in one 39-mer half than in the other, reflecting the presence of a longitudinal compression between the primed and the committed conformations.

**Figure 3.**
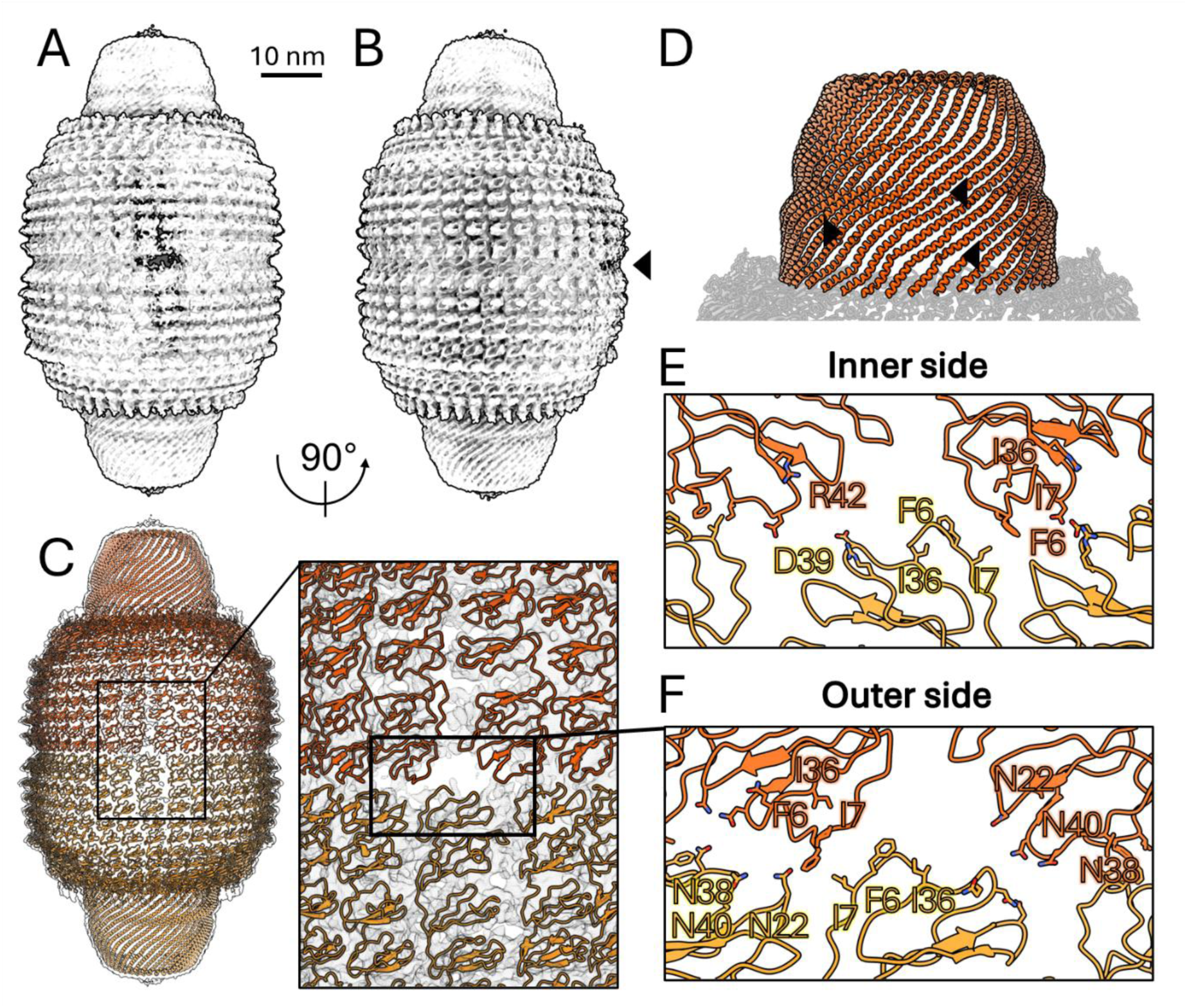
**Volume and structure of the vault in committed conformation**. A, B) Cryo-EM density map of the vault in committed conformation (contour level 0.35). The black arrow indicates the region of distortion at the waist. C) Atomic structure of the vault in committed conformation, superimposed onto the cryo-EM map. (Inset) A detail of the atomic structure of the vault in committed conformation, with the density map shown as black mesh (contour level 0.35). D) Structure of the helix bundle in the cap-helix. The black arrows indicate the kinks in the helices induced by the conformational change. E, F) Atomic model of the vault’s waist, shown from the inner side (E) and outer side (F). Cartoon representation of the vault in committed conformation, with residues at the interface of the two 39-mer MVP halves shown as sticks, illustrating the loss of contacts in the region of greater distortion. Hydrophobic patch residues (Phe6, Ile7 and Ile38), the polar residues of the polar triad (Asn22, Gln38 and Asn40), as well as the charged residues (Asp39 and Arg42) are shown as sticks.

The atomic model of the committed vault was obtained through symmetry relaxation and molecular dynamics flexible fitting (MDFF)^46,47^ of the symmetric higher-resolution primed structure into the committed density map (**Figure 3C**). Besides the extensive distortion of the waist observed in the committed conformation (**Figure 3B**), we noted further structural deformations in the cap-helix, where several kinks in the α-helices were observed (**Figure 3D**). Additionally, we performed a local refinement of the midsection of the vault aimed at improving the resolution of the R1-R2 domains at the waist, which resulted in a local refinement map at resolution of 6.06 Å (**Supplementary Figure 10B**).

The heterogeneity of the particles used in the reconstruction of the map resulted in a markedly lower resolution in the symmetry-mismatched component at the waist, leading to a low-confidence model for the distorted repeat domains. Nonetheless, the curvature of the vault in the region of greatest displacement visibly forced the two 39-mer halves apart and determined an overall loss of lateral contacts along R1-R6 domains and at the level of opposing R1 domains. Specifically, the interaction between two antiparallel R1 domains was lost due the disruption of the inter-chain salt bridge formed between Asp39 and Arg42 and the absence of the tight interaction between residues Phe6, Ile7 and Ile38 of two opposing R1 domains (**Figure 3E**). Furthermore, in two instances, we observed a lack of polar interactions between the opposing polar triads (Asn22, Gln38 and Asn40) at the waist interface (**Figure 3F**).

Next, the structure of the vault in the committed conformation was closely compared to that of the primed conformation. Most noticeably, the symmetry-mismatched component in the committed conformation stretched the vault’s waist. Compared to the diameter in the primed conformation (35 nm × 35 nm), the committed conformation assumed a more elliptical shape, with a diameter of 37 nm × 34 nm. (**Figures 4A** and **4B**). Moreover, the diameter of the helix bundle in the cap helix is smaller than in the primed conformation by up to about 1 nm (**Figures 4B** and **4C**). This distortion was markedly more prominent in one 39-mer half than the other, indicating that relaxation at the waist, associated with the loss of N-terminal contacts between the two halves, propagated longitudinally and culminated in changes in the cap-helix (**Figure 4A**). This transition was also captured by the 3DVA performed on the whole particle stack (**Supplementary Movie 1** and **Supplementary Figure 4**).

**Figure 4.**
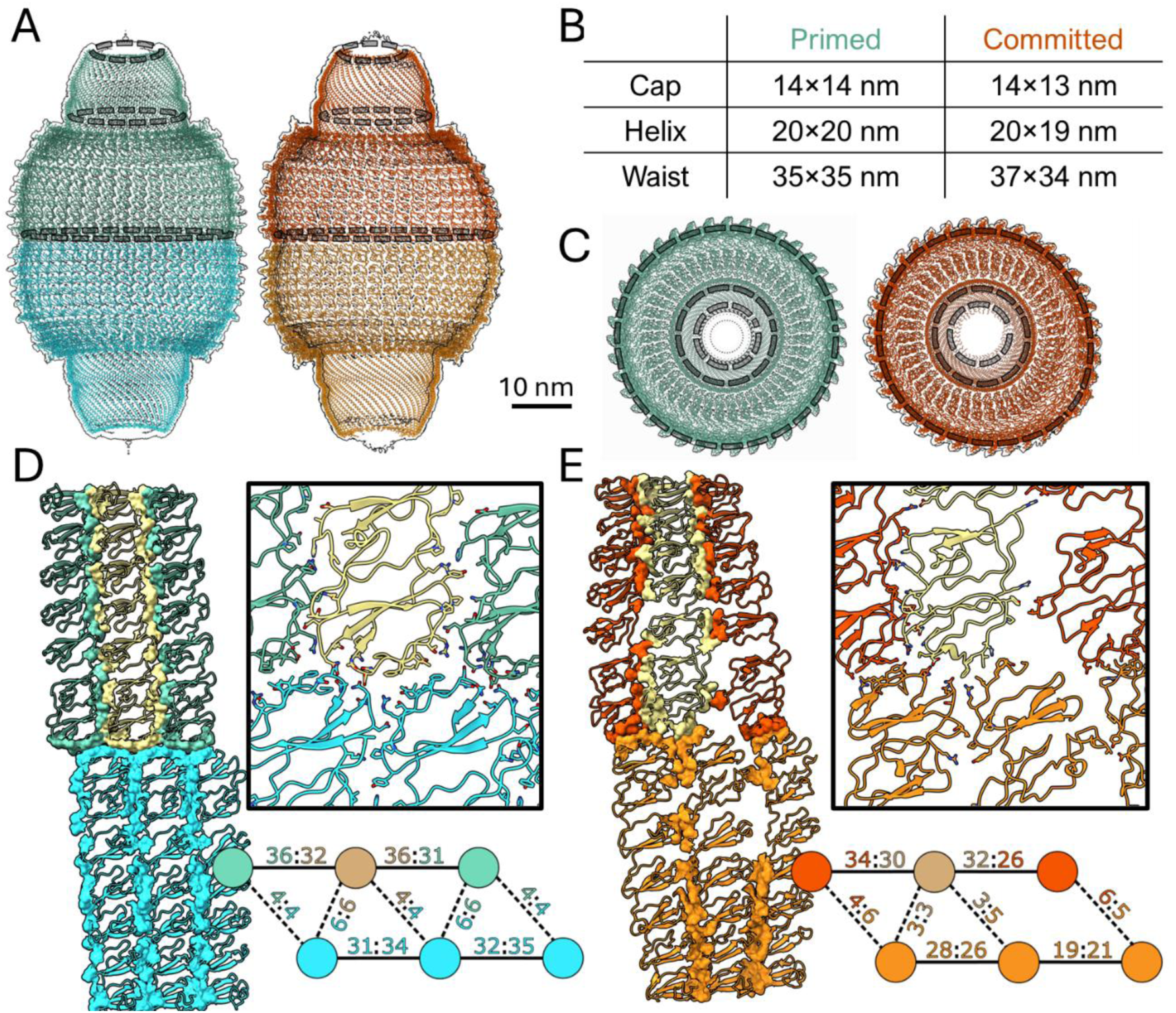
**Structural comparison between vault in primed and committed conformations**. A) Clipped representation of vault in primed and committed conformations (left and right, respectively). Cartoon representation of the backbone and map shown as a surface. The dashed circle indicates the cross sections used to calculate the diameter B) Table comparing the diameter of the two structures (see Methods) at different cross-sections (shown in A and C as dashed circles). C) Top view of the upper 39-mer half of the vault in primed and committed conformations (left and right, respectively). In (A, C), contour levels are set at 0.9 and 0.4 for primed and committed, respectively. D, E) Left panel, lateral view of six MVP chains (residues 1-379) in the primed (D) and committed (E) vault structures, respectively. Cartoon representation of the model with visible surface of the residues burying at least 11Å^2^ of area at the interface of different MVP chains. The top-right panels show a detailed view of the six chains with contacting residues as sticks. The bottom-right panels show a schematic representation quantifying the number of contacting residues among the six chains (between residues 1-379), illustrating the quaternary organization of the vault protein. Each polypeptide chain is shown as a coloured circle. Black solid lines connect chains within the same 39-mer half of the vault, whereas dotted lines indicate inter-subunit contacts that bridge the two 39-mer halves. The numbers on each connection (separated by a colon) identify the number of residues from each chain that participate in these intermolecular contacts.

Analysis of the contacts between R1-R7 domains in the primed conformation showed tightly packed repeated interfaces between chains. Each MVP monomer interacted with four other chains, two along the waist of the vault and two opposing on either side (**Figure 4D**). The waist interactions are stabilized by hydrophobic patches, whereas a network of polar interactions zips up the opposing chains. In the committed conformation, by contrast, MVP chains located in the most distorted region of the waist showed an overall loss in the number of lateral contacts and a complete lack of interactions between two opposing chains (**Figure 4E**). This ultimately resulted in the formation of a gap between the two antiparallel 39-mer halves and of longitudinal pores along the interface of parallel MVP subunits (**Figures 3C** and **4E**).

### The half vault

The presence of a sizeable population of 39-mer half vaults on the micrographs allowed us to reconstruct a low-resolution map of the disassembled half particle at resolution of 9.89 Å (**Supplementary Figure 11**). The reconstructed volume indicated a clear deviation from the regular conformation assumed by the halves in the whole vault. The waist appears distorted, and the heterogeneity of the particles resulted in a lower-density region in the map, corresponding to the R1-R6 domains. This suggests a higher degree of flexibility in this region, in line with the fewer lateral contacts present. The loss of structural constraints at the waist clearly affects the curvature of the six initial repeat domains in the half vaults, which closely resemble the 39-mer halves of the vault in committed conformation.

### Analysis of vault dynamics

To obtain a detailed view of the conformational dynamics of the vault complex, we performed classical all-atom molecular dynamics (AA-MD) and coarse-grained molecular dynamics (CG-MD) simulations, focusing on the primed, committed, and half-vault states. Starting from the distinct cryo-EM structures, we built atomistic models to characterize the dynamic interactions of MVP subunits.

In our simulations, we included all MVP subunits (residues 1–814), fully solvated in water with Na⁺ and Cl⁻ ions (**Figure 5A**). We performed two replicates of AA-MD simulations for each initial conformation, each running for 250 ns, yielding a cumulative simulation time of 1 μs. These AA-MD simulations involved systems of ∼18.4 million atoms (**Supplementary Table 2**), highlighting the feasibility of studying such a large biological assembly, thanks to advances in supercomputing resources^48^. The AA-MD simulations revealed that both conformations largely maintained their overall structural integrity throughout the simulation, as illustrated by representative snapshots from the MD trajectories (**Figure 5B**). Nevertheless, the interfaces at the level of the symmetry-mismatched component in the committed conformation displayed greater variability, with the prominent rupture at the waist showing attempts to close but ultimately forming fewer stable contacts compared to the primed conformation.

**Figure 5.**
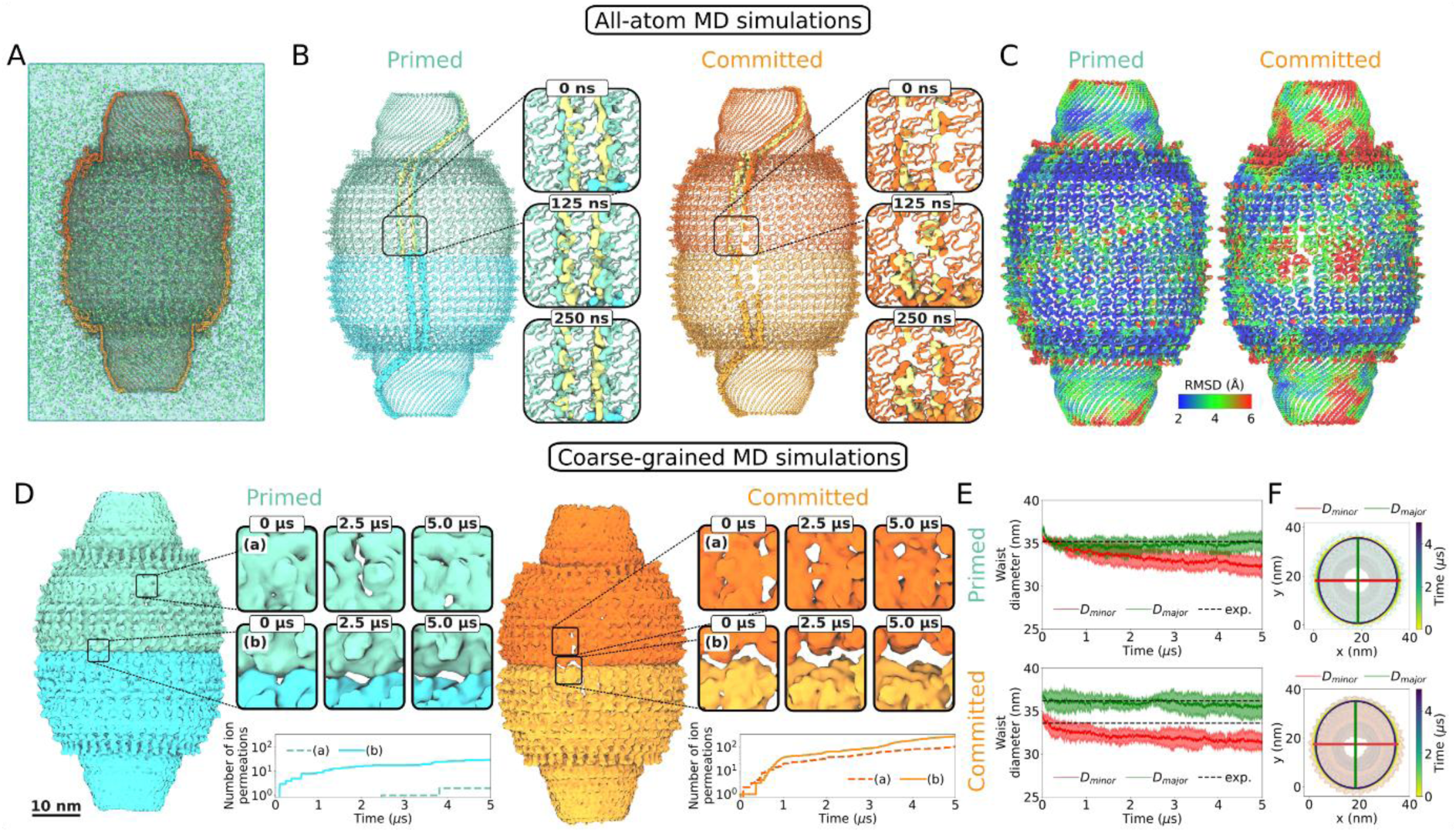
**All-atom (A–C) and coarse-grained (D–F) molecular dynamics simulations of the vault in primed and committed conformations**. A) Cutaway view of the all-atom MD simulation box containing the human vault particle in the committed conformation (orange), solvated in water (cyan surface) with Na⁺ (purple) and Cl⁻ (green) ions. B) All-atom MD simulations of the vault in primed (left) and committed (right) conformations. Cartoon representation of the initial atomic models, with surfaces highlighting inter-MVP contacts. Zoomed-in views show representative snapshots from the MD trajectory. C) RMSD over the last 200 ns of simulation, mapped onto the initial atomic structures of the vault in primed (left) and committed (right) conformations. Structures are aligned to the Cα atoms of residues with RMSF <3.5 Å. D) Coarse-grained MD simulations of the vault in primed (left) and committed (right) conformations. The coarse-grained structures are shown as surfaces (contour level 1.3). Zoomed-in views highlight the dynamic behaviour at MVP interfaces (a) and at the waist in the interface between vault halves (b) creating pores. Cumulative number of ion permeations through zoomed-in regions (a) and (b) in the vault shell (bottom), measured by tracking ions entering and exiting the vault through pores in each region. E) Time series of the vault’s waist diameter in the primed (top) and committed (bottom) conformations. Diameters were obtained by fitting an ellipse to the backbone bead positions of residues D39, with major and minor diameters shown in green and red, respectively. Mean values (solid lines) and two standard deviations (shaded regions) from three simulation replicates are plotted. Dashed lines indicate diameters from the atomic model. F) 2D representation of the fitted ellipses for the vault’s waist diameter over time in the primed (top) and committed (bottom) conformations. Half-vaults are shown in transparency for reference. Each ellipse is constructed using the major and minor diameters fitted to the backbone bead positions of residues D39. The colour scale represents the time point at which each ellipse was fitted, with one ellipse plotted every 150 ns.

The primed conformation showed a higher stability in AA-MD simulations compared to the committed conformation. To visualize the dynamic regions of both conformations, we mapped the RMSD from the starting model over the last 200 ns of AA-MD onto their initial atomic structures, aligning them to the Cα atoms of residues with root-mean-square fluctuation (RMSF) <3.5 Å (**Figure 5C**). The primed conformation exhibited fewer overall fluctuations, while the committed conformation showed increased flexibility, particularly at the waist, cap, and shoulder regions. This increased flexibility was consistent with the lower resolution of the cryo-EM model and the stretching-compression motions observed in the 3DVA. Despite these fluctuations, the AA-MD simulations accurately captured the general shape of the human vault particle with remarkable agreement, as shown by particle diameter measurements at the waist, helix, and cap regions (**Supplementary Figure 12A**). Although AA-MD simulations provide atomic-level detail, their accessible timescales remain relatively short compared to those needed to capture large conformational changes. CG-MD simulations address this limitation by enabling the exploration of larger-scale dynamics over extended timescales. For the human vault particle, the system size was reduced to ∼1.6 million beads, allowing us to perform 5 μs of simulation for three replicates per initial configuration to further assess the vault’s flexibility. Representative coarse-grained structures of the primed and committed conformations revealed the formation of dynamic horizontal pores at the waist and vertical pores between MVP subunits (**Figure 5D** and **Supplementary Movie 2**). Quantification of cumulative ion permeation through specific regions at the vault shell shows that regions with visually larger pores allow greater ion permeation (see bottom panels in **Figure 5D** and Methods).

The vault particle exhibited remarkable flexibility in CG-MD simulations, particularly at the waist. To quantify this dynamic behaviour, we tracked changes in waist diameter over the course of the simulation by fitting an ellipse to the backbone bead positions of residues Asp39, with the major and minor diameters shown in green and red, respectively (**Figure 5E**). The time series data revealed substantial flexibility at the waist, with good agreement between the major diameter of the fitted ellipses and the cryo-EM atomic model. These fluctuations suggested the presence of a general breathing motion. A two-dimensional representation of the fitted ellipses over time illustrated how the vault gradually adopted a more ellipsoidal shape as the CG-MD simulations progressed (**Figure 5F**). This ellipsoidal shape in both the primed and committed conformations associated with the expansion of the waist pores.

We also measured the diameters of the cap and helix regions for both vault configurations (**Supplementary Figure 13A**). Our CG-MD simulations systematically underestimated the cap diameter, likely due to the absence of the intrinsically disordered C-terminal regions of MVP. By contrast, the helix diameter closely matched the cryo-EM models. Both the cap and helix regions displayed smaller diameter fluctuations than the waist, supporting the hypothesis that the vault’s greater flexibility at the waist is closely linked to its opening mechanism.

To further characterize vault flexibility, we analyzed individual half-vaults starting from both the primed and committed conformations, comparing AA-MD and CG-MD simulations with experimental cryo-EM data. Given the larger structural fluctuations observed in the committed state, we focused on this conformation in the main text (**Figure 6**), while results from the primed half-vault are presented in the supplementary information (**Supplementary Figure 14**). AA-MD simulations of the half-vault system, comprising ∼10.7 million atoms (**Figure 6A**), revealed greater structural changes, particularly at the waist, compared to the full vault particle. This is reflected in the RMSD from the last 200 ns of AA-MD simulations (**Figure 6B-left**), where higher RMSD values indicate increased flexibility and more pronounced conformational changes. Interestingly, R1-R6 regions show large variations in RMSD, consistent with the large distortions in the experimental cryo-EM map of the 39-mer halves (**Figure 6E-bottom**). The RMSD of CG-MD simulations over the last 4.5 μs showed not only large instability at the waist but also increased flexibility at the cap region (**Figure 6B-right**). Tracking the waist diameter revealed greater fluctuations and larger amplitude breathing movements compared to the full vault (**Figure 6C**), underscoring the enhanced flexibility of the half-vault.

**Figure 6.**
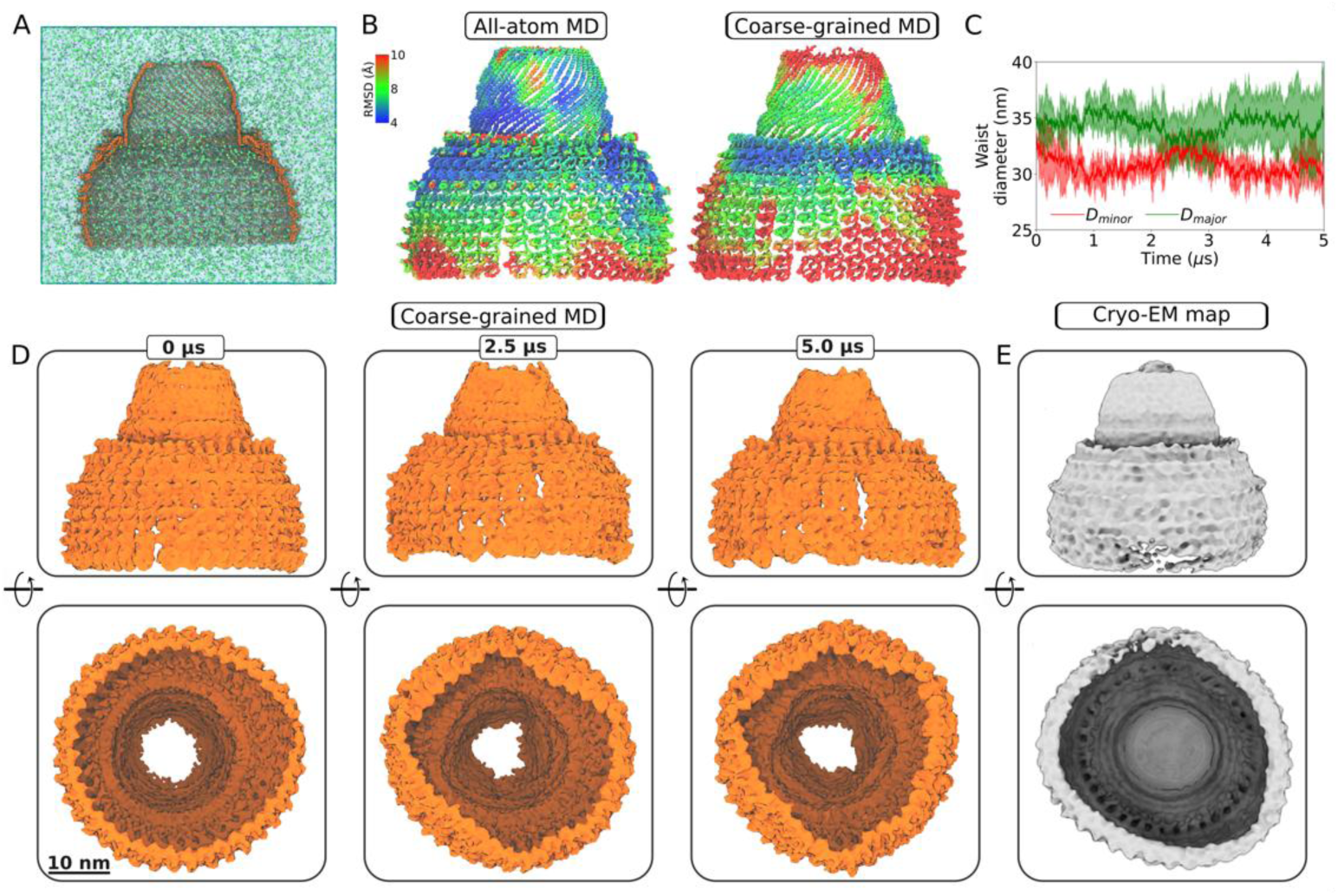
**All-atom (A–B) and coarse-grained (C–D) molecular dynamics simulations of the half-vault in the committed conformation**. A) Cutaway view of the all-atom MD simulation box containing half of the human vault particle in the committed conformation (orange), solvated in water (cyan surface) with Na⁺ (purple) and Cl⁻ (green) ions. B) Left panel, RMSD over the last 200 ns of simulation, mapped onto the initial atomic structure of the half-vault in the committed conformation. Structures are aligned to the Cα atoms of residues with RMSF <3.5 Å. Right panel, RMSD over the last 4.5 μs of simulation, mapped onto the initial coarse-grained structure of the half-vault in the committed conformation. Structures are aligned to the backbone beads of residues with RMSF <3.5 Å. C) Time series of the half vault’s waist diameter in the committed conformation. Diameters are obtained by fitting an ellipse to the backbone bead positions of residues D39, with major and minor diameters shown in green and red, respectively. Mean values (solid lines) and two standard deviations (shaded regions) from three simulation replicates are plotted. D) Coarse-grained MD simulations of the half-vault in the committed conformation. Representative snapshots from the MD trajectory are shown from a lateral view (top) and an internal bottom-up view (bottom). The coarse-grained structures are represented as surfaces (contour level 1.3). E) Lateral (top) and internal bottom-up (bottom) views of the non-uniform cryo-EM refinement of the 39-mer half-vault, reconstructed from 3,069 particles (contour level 0.3).

Representative snapshots from the CG-MD trajectory, viewed laterally and from an internal bottom-up perspective, revealed prominent structural deformations, including the formation of large vertical pores and inward movement of the MVP N-terminal regions at the waist (Figure 6D). The low-resolution cryo-EM map (**Figure 6E**) further suggested significant conformational flexibility, with the waist shape closely resembling the deformations observed in the CG-MD simulations. In contrast, simulations starting from the primed half-vault showed smaller variations in waist diameter and overall lower RMSD values (**Supplementary Figure 14**).

## Discussion

Recent methodological developments in cryo-EM analysis facilitate the study of continuous flexibility and discrete heterogeneity of large molecular assemblies^49^. In this work, we harness these advances to perform a comprehensive structural analysis of the human vault particle at atomic resolution. We investigated the molecular determinants driving the transition of the vault from a symmetric conformation – primed – to a state in which the symmetry is lost – committed.

Cryo-EM reconstruction of the human vault particle revealed a structure mirroring its murine counterpart. Atomic-level details of the structure showed an intricate network of interactions that stabilize each monomer. Focusing on the interface between the two halves in the primed conformation, we identified the presence of a hydrophobic patch and two distinct sets of polar residues that stabilize the two anti-parallel 39-mer halves, which are crucial for the vault overall stability. Conversely, the committed conformation, a presumed on-path intermediate in the vault opening process, showed a discernible distortion at the waist diameter. Differential analysis of the two conformations at the level of monomer-monomer contacts revealed the transition from primed to committed as an overall loss of contacts at the waist, specifically involving the disruption of one hydrophobic patch and loss of interactions between two polar triads and a salt bridge facing opposing R1 domains at the waist. This suggested that the increased structural heterogeneity observed in the committed conformation is driven by weakened interactions between opposing antiparallel R1 domains, compromising the stability of the assembly and potentially facilitating the subsequent disassembly of the vault.

These two alternative conformations of the vault, further analyzed by MD simulations, emerged to be distinct states. In the MD simulation trajectories, the vault underwent significant breathing motions in solution, yet the particle did not transition between the two conformations, nor did the two halves initiate the opening in the absence of external stimuli. Nonetheless, MD simulations revealed that the vault is dynamic. The architecture of the vault is the result of a network of weak interactions than can break and form due to thermal fluctuations, resulting in the opening of further pores on the interfaces besides those present in the initial cryo-EM structure during the simulations. These pores could serve as a mechanism for pressure release and passive diffusion of ions and small molecules.

The committed conformation of the vault exhibits greater structural variability, as reflected by higher RMSD values in MD simulations, consistent with the higher structural heterogeneity observed in cryo-EM for this conformation. MD simulations revealed the formation of indentations, particularly at the waist region, a feature that has also been directly imaged in cells^50^. Moreover, half-vault particles showed enhanced fluctuations at the waist, emphasizing the critical role of inter-half contacts in stabilizing the vault’s overall architecture. Notably, MD simulations starting from the primed half-vault cryo-EM model displayed less structural dynamics compared to those initiated from the committed half-vault model.

Taken together, our results suggest that the two identified human vault structures are bona fide representations of alternative conformational states of the vault particle, with the committed conformation being an intermediate in the pathway towards opening. Higher flexibility at the interface between the two 39-mer halves observed in the simulations should facilitate a breakup at the waist. In the EM structures, the local disengagement of the N-terminal domain through the loss of both polar and hydrophobic interactions at the interface between the two halves (**Figure 3E** and **3F**) is connected to a change in diameter of the vault’s waist. The presence of 39-mer halves found in the SAXS analysis and identified at low resolution by cryo-EM further pointed to the presence of an equilibrium between the two states, supporting the model of vault opening and closing at the waist.

The similarity of the conformations solved in our study to those initially observed in the murine vault points to the generality of the proposed dissociation initiation mechanism^38^. In addition, providing a high-resolution structure of the vault in the committed conformation and examining its flexibility with MD simulations allowed us to build upon the proposed mechanistic model for the opening of the vault and to pinpoint the structural elements relevant in intermediate conformations.

As seen in other human capsid protein assemblies involved in cellular and inter-cellular transport, such as ferritin^51^ or the activity-regulated cytoskeleton-associated (arc) protein^52^, the disassembly process of the vault, might have profound implications for its physiological role as carrier, both intracellularly and, as suggested by recent findings, also extracellularly^53,54^. Furthermore, the vaults attractiveness as self-assembling human nanoparticles, simultaneously useful for both targeted delivery and encapsulation^55–58^, emphasizes the importance of understanding their dynamics, as well as elucidating the molecular mechanisms underlying their opening and closing. Nevertheless, while our study focused on the molecular nature of vault’s unique morphology and structural flexibility, further studies will be required to fully elucidate the disassembly process in its completeness.

## MATERIALS and METHODS

### MVP expression and purification

The human MVP gene (GenBank: BC015623.2) was cloned in the expression vector pGAPZB (Invitrogen) as already described in^41^. Briefly, the plasmid was amplified in *E. coli* strain DH5α growth in low salt LB media (10 g/L Bacto-tryptone, 5 g/L yeast extract and 5 g/L NaCl, pH 7.0) supplemented with Zeocin (50 µg/mL), purified with silica spin-columns (Qiagen) and linearized with EcoRI and KpnII (NEB). The PEP4 protease deficient strain of *K. phaffii* SMD1168 (his4, ura3, pep4:URA3), growth for 24 hours in YPD media (20 g/L Bacto-peptone, 10 g/L yeast extract, pH 7.0) was electroporated using a Gene Pulser II electroporator (Bio-Rad) with a single pulse at 1.5 kV, 400 Ω and 25 μF and plated on YPD agar plates containing 1 M sorbitol and 100 µg/mL zeocin. Single colonies were further screened by colony-PCR amplification with Taq polymerase (ThermoFisher Scientific) and by western blot, developed with anti-MVP antibody produced in rabbit (SAB5700906 – Sigma-Aldrich).

Selected clones were inoculated in YPD supplemented with zeocin (100 µg/mL), growth at 30 °C for 24 hours and stored at-80 °C, in presence of 25% v/v glycerol. The stocks were thawed and grown in YPD at 30 °C for 72 hours in agitation, harvested, washed twice with 50 mM Tris-HCl, 150 mM NaCl, pH 8.0 in presence of 3 mM of dithiothreitol (DTT) and then lysed. Previous addition of 1 mL of lysis buffer per gram of cellular pellet (50 mM Tris-HCl, 150 mM NaCl, 1mM TCEP, pH 8.0), plus 1 U/mL of DNase I (VWR), 1 U/mL RNase T1 (ThermoFisher Scientific), 1 U/mL RNase A (Sigma), 1 mM ATP, 1 mM PMSF, protein inhibitor cocktail (Roche), 1 mM EDTA and 5v/v glycerol, the lysis was performed on ice by alternate cycles of vigorous shaking in presence of 25 % w/v of glass beads and centrifugation at 10,000 x *g*. After removal of the insoluble fraction by centrifugation at 20,000 x *g* for 1 hour at 10 °C, the lysate was loaded onto a cushioned discontinuous density gradient of sucrose composed of three levels (20% w/v, 45% w/v and 60% w/v) and applied to ultracentrifugation at 100,000 x *g* for 2 hours at 10 °C, The sample was recovered from the 45% level, diluted to a final concentration of sucrose below 10% and incubated with 1 U/mL RNase T, 1 U/mL RNase A, 1 mM ATP and 1 mM CaCl_2_ for 30 minutes at room temperature and centrifuged again at 16,000 x *g* to remove aggregates.

The sample was then loaded into a Sephacryl S-500 HR column 16/60 (Cytiva) previously equilibrated with SEC buffer (25 mM HEPES, 150 mM NaCl, 1mM TCEP, pH 7.5), and eluted at 2.4 mL/min (elution peak between 140 mL and 180 mL). The fractions were analysed by SDS-PAGE stained with colloidal Coomassie InstantBlue (abcam) and by transferring the sample on a PVDF membrane, incubated with anti-MVP (SAB5700906 – Sigma-Aldrich), and developed with mouse anti-rabbit IgG-HRP, followed by chemiluminescence imaging (UVITEC). Finally, the sample was concentrated with ultrafiltration centrifugal devices with 100 kDa cut-off membrane (Millipore); the protein concentration was estimated spectrophotometrically using the theoretical extinction coefficient at 280 nm of 5×10^6^ × M^−1^ × cm^−1^ obtained from Expasy servers^59^.

### Protein characterization

*Size-exclusion chromatography coupled to multi-angle light scattering (SEC-MALS).* SEC-MALS was used to calculate the molecular mass of the sample. The Dawn Heleos II MALS detector (Wyatt), coupled to a HPLC system Alliance e2695 (Waters) was used to read the intensity of the scattered light using 8 angular detectors. The sample was filtered through 0.2 μm centrifugal filter units (Millipore), injected into a Shodex KW405-4F column (Shodex) and eluted at 0.33 mL/min in SEC buffer. The molecular weight of the eluted peak was calculated integrating the signal of refractive index detector RI-501 (Shodex) and the MALS detector with the software ASTRA (version 7.0) (Wyatt).

*Small-angle X-ray scattering (SAXS).* SAXS data were collected at the European Synchrotron Radiation Facility (ESRF, Grenoble, France) on the BM29 BioSAXS beamline. Scattered X-rays at a wavelength of 0.992 Å (E = 12.5 keV) were recorded using a PILATUS3 2M (Dectris) detector in the SAXS region (q = 0.025–6 nm⁻¹). Two samples at a concentration of 0.5 mg/mL and 0.3 mg/mL were measured in buffer containing 25 mM HEPES, 150 mM NaCl, and 1 mM TCEP (pH 7.5). All samples were centrifuged at 20,000 × *g* for 10 minutes at 4°C prior analysis. Diluted samples were injected directly using the sample changer at 20°C. For each sample measurement, ten X-ray scattering measurements with 1-second exposure times were collected. These ten analogous signal frames were averaged, and buffer scattering was subtracted from the sample data.

Pair distance distribution functions were calculated with the GNOM program that is part of the ATSAS program suite^60^ for small-angle scattering data analysis (version 3.0.4). Fitting of single atomic models to the SAXS scattering curve was performed with PepsiSAXS^61^ (version 3.0), which employs an adaptive multipole expansion approach to efficiently and accurately compute scattering intensities. Multi-component fitting was instead done with OLIGOMER program that is part of ATSAS.

### Cryo-EM sample preparation and imaging

The sample was concentrated to 250 nM and biotinylated with 10 molar excess of biotinylation reagent (NHS-PEG_12_-biotin) for 30 minutes at room temperature. The reaction was quenched with 1 mM Tris HCl pH 7.5 and excess of unbound reagent was discarded by 200-fold dilution with sample buffer (25 mM HEPES, 150 mM NaCl, 5 mM CaCl_2_, 1 mM TCEP, pH 7.5) followed by ultra-filtration, final sample concentration was estimated to ∼ 50 nM. Gold film grids functionalized with 2D streptavidin crystals (SAG), prepared and stored according to the procedure already described in^62,63^ were rehydrated, equilibrated with sample buffer and incubated in sealed plate at 4°C with 5 µL droplets of biotinylated samples. After 15 min, wells were unsealed, and samples were homogenized by pipetting in-and-out 2 µL. After further 15 min, Grids were lifted up from plate pedestals with 100 µL sample buffer, recovered, further washed on 100 µL sample buffer, plunged frozen using a Vitrobot Mark IV (ThermoFisher Scientific) with 5 seconds blot time and kept in liquid nitrogen until imaging.

Data collection was performed on a Titan Krios microscope (ThermoFisher Scientific) at 300 keV, equipped with a CS-corrector, Gatan K3 electron detector and a GIF bioquantum energy filter, with calibrated pixel size of 0.8416 Å × pix^−1^, spherical aberration of 0.01 mm and electron dose rate of 50.15 e^−^/Å^2^.

### Cryo-EM Single particle analysis

A dataset of 6,895 movies was recorded and initially processed in RELION^64^ (version 3.1). The streptavidin crystal lattice of the SAG was subtracted from the micrographs in Fourier space with a python script already described in^63^ and available at https://github.com/NilsMarechal/SAGsub. Subtracted micrographs were then imported and further processed into cryoSPARC (version 4.5.3)^43^. Contrast transfer function (CTF) was estimated for all the micrographs and used to exclude the ones with CTF > 7 Å. A selection of manually picked particles (∼ 250 particles) was used to generate 2D classes for template-based picking. To remove junk particles, contamination and artifacts, 63,814 particles (box size of 1024 × 1024 pixels at pixel size of 0.8416 Å) were selected for 2D classification followed by ab initio 3D reconstruction. This allowed to differentiate picked particles between vaults and half vaults. Rounds of 2D classification, heterogeneous refinement, 3D classification and homogeneous refinement were employed to further curate the stack of particles and remove partially disassembled vaults artifacts using down-sampled particles in Fourier space to 550 × 550 pixels box size. An initial homogeneous refinement at GS-FSC resolution of 4.28 Å was used to perform 3D classification of the whole vault. Filter resolution at 10 Å was applied to classify the particle stack into the two conformations discussed in our work: primed and committed. Non-uniform refinement was performed after re-extracting the particles with 1024 × 1024 pixels box size for both stacks with CTF corrections (beam tilt and trefoil), minimizing per-particle defocus and scale, with 5 extra passages after GS-FSC resolution stopped improving. The final map for the primed conformation at GS-FSC resolution of 3.09 Å was obtained imposing dihedral 39-fold symmetry (D39). Applying no symmetry restrain to the refinement (C1) resulted in a similar map, lacking symmetry-mismatched components, albeit with an overall lower resolution of 4.78 Å. D39 symmetry relaxation without pose marginalization was used to refine the map of the committed conformation at GS-FSC resolution of 4.45 Å, a comparable resolution of 4.51 Å was obtained by refining the map in the absence of any imposed symmetry (C1) as the former map resulted in more legible density at the waist this was selected for the atomic reconstruction. To improve the resolution of the map in the regions of lower density, local refinements were performed with masks generated around the waist. The maps were sharpened with the negative B-factor (Å^2^) estimated after the refinement and all GS-FSC resolutions were determined using the gold-standard FSC at 0.143, with FSC curves adjusted for a tight mask in cryoSPARC^43^ (version 4.5.3).

### Model building

The molecular model of the vault in primed conformation was built starting from the structure of a MVP monomer (residues 1-814) generated by homology to the structure of crystallized MVP (PDB: 4HL8)^10^ using MODELLER^65^ (10.5). Coot^66^ (0.8.9) and ChimeraX^67^ (version 1.8) were employed to rebuild missing loops and to perform initial fitting to the main map taking in consideration favoured Ramachandran dihedral angles, rotamers and clashes. The map resulting from the local refinement of the waist was used to build the N-terminal domains R1-R2. Non-crystallographic symmetry of the whole map was identified and applied to reconstruct an initial model of the full 78-mer assembly using PHENIX^68^ (1.21). Cycles of improvement of the fit between the map and the atomic model of a pentameric unit of MVP were performed using ISOLDE^47^ in ChimeraX^67^ (version 1.8). Symmetry expansion according to the D39 point group symmetry of the central MVP monomer of the pentamer and subsequent real-space refinement of the 78-mer MVP was performed using PHENIX^68^ (version 1.21).

The model of the vault in committed conformation was obtained through cycles of improvement of the fit between the atomic structure of the vault in primed conformation with the maps of the vault in committed conformation using PHENIX real-space refinement and Molecular Dynamics Flexible Fitting (MDFF) in ISOLDE^47^. For MDFF, the weighting between the map and the model parameters was carefully adjusted to avoid overfitting. Initial energy minimization was followed by a two-step molecular dynamics procedure; a first short step at low weighting (0.1) followed by a second step at higher weighting (0.3), for the whole all-atom model and subsequently only for the chains in the symmetry-mismatched component (using the map resulting from the local refinement of the waist). The full-length models were used to perform the molecular dynamics simulations. Afterwards, the following segments (residues 1-2, 429-449, 608-619), where the low resolution of the map prevented an accurate modelling, were trimmed from both models, which were thereby subjected to 20 macro-cycles of global real-space refinement^69^ in PHENIX (version 1.21).

### Molecular Dynamics

We performed all-atom and coarse-grained MD simulations of vault particles and half vault particles in primed and committed conformations. In all cases, we started from the atomistic models of the corresponding high-resolution cryo-EM structures solved in this work. All the MD simulations reported in this work were performed in GROMACS 2024^70^ and are listed in **Supplementary Table 2**. The MD simulation setup, MD trajectory analysis and visualization are described below.

*All-atom MD simulations.* The atomistic structures of vault and half-vault particles were used as input for all-atom MD simulations. The AMBER99SB-ILDN^71^ force field was applied, and the systems were solvated with TIP3P water and 150 mM NaCl. Energy minimization was performed using the steepest-descent algorithm until the maximum force was <1000 kJ·mol⁻¹·nm⁻¹. Equilibration proceeded in multiple stages with position restraints (force constant: 1000 kJ·mol⁻¹·nm⁻²) applied to protein heavy atoms. All simulations employed the Verlet neighbour search algorithm (neighbour list cutoff: 1.2 nm, update frequency: 20 timesteps). Lennard-Jones interactions were truncated at 1.2 nm. Long range electrostatics were treated with Particle Mesh Ewald and truncated at 1.2 nm. (i) NVT equilibration for 375 ps with a 1 fs timestep, using the Berendsen thermostat^72^ (T = 310 K, τ = 0.01 ps). (ii) NPT equilibration for 1 ns with a 2 fs timestep, using isotropic pressure coupling via the Berendsen barostat^72^ (P = 1 bar, τ = 2 ps) and switching to the v-rescale thermostat^73^ (T = 310 K, τ = 0.1 ps). (iii) NPT equilibration for 100 ns with a 2 fs timestep, using the v-rescale thermostat (T = 310 K, τ = 0.1 ps) and the c-rescale barostat^74^ (P = 1 bar, τ = 2 ps). (iv) Final NPT equilibration for 20 ns under the same conditions as (iii) but with the position restraint force constant reduced to 10 kJ·mol⁻¹·nm⁻². Two production runs of 0.25 μs per initial structure were performed following the protocol in (iii), without position restraints on protein heavy atoms.

*Coarse-grained MD simulations.* The atomistic structures of vault and half-vault particles were used as input to generate coarse-grained (CG) Martini protein models. Each MVP chain was coarse-grained individually using the martinize.py script^75^, with default protonation states and secondary structure restraints assigned via DSSP^76^. All CG simulations employed the Martini 2.2 force field^77^ alongside the ElNeDyn2.2 protein force field^78^. Given the known overestimation of nonbonded interactions^79^ in Martini 2.2, we used an α parameter of 0.85 to scale protein-protein interactions relative to protein-solvent interactions^80^. To maintain protein stiffness, an elastic network with bond cutoffs of 0.5 and 0.9 nm was applied to each MVP chain with elastic bond force constant of 1000 kJ·mol⁻¹·nm⁻².

The systems were solvated with coarse-grained water containing 10% anti-freeze WF particles and 150 mM NaCl, with excess ions ensuring charge neutrality. Energy minimization was performed using the steepest-descent algorithm and a maximum force < 1000 kJ·mol⁻¹·nm⁻^1^. Equilibration proceeded in three stages with position restraints (force constant: 1000 kJ·mol⁻¹·nm⁻²) applied to protein backbone beads: (i) NVT equilibration for 1.0 ns with a 5 fs timestep, using the Berendsen thermostat^72^ (T = 310 K, τ = 1 ps). (ii) NPT equilibration for 2.5 ns with a 5 fs timestep, applying isotropic pressure coupling via the Berendsen barostat^72^ (P = 1 bar, τ = 12 ps) and switching to the v-rescale thermostat^73^ (T = 310 K, τ = 1 ps). (iii) Final NPT equilibration for 300 ns with a 15 fs timestep, using the v-rescale thermostat (T = 310 K, τ = 1 ps) and the c-rescale barostat^74^ (P = 1 bar, τ = 12 ps). Three production runs per initial structure of 5 μs each were conducted using the same protocol as (iii) but without position restraints on backbone beads. All simulations employed the Verlet neighbour search algorithm (neighbour list cutoff: 1.4 nm, update frequency: 20 timesteps). Lennard-Jones and Coulomb interactions were truncated at 1.2 nm using the Verlet-shift potential modifier and reaction-field electrostatics. All MD simulations were performed at the AMD-based supercomputer VIPER of the Max Planck Society, operated at the Max Planck Computing and Data Facility in Garching.

Visual Molecular Dynamics (VMD)^81^ was used for trajectory visualization, while MDAnalysis^82^ was employed for trajectory analysis, RMSD and diameter measurements.

*Ion permeation analysis.* To quantify the cumulative crossing of ions through specific regions in the vault shell, we analysed the coarse-grained MD trajectories of the vault particle using MDAnalysis and custom Python scripts. A region was defined by the convex hull of six neighbouring R domains. A vector **A** between the centre of mass (CoM) of the full vault shell and the CoM of the convex hull defined the main axis of a cylindrical volume. This volume, with a radius =3 nm, was used to spatially constrain the analysis to ions passing directly through the selected R domains. At each frame, we determined which ions (Na⁺ or Cl⁻) are within the cylindrical volume. Furthermore, we define the vector **B** between the ion position and the CoM of the convex hull. For the cases where, |𝐀 ⋅ 𝐁| > 1 nm, the ion was considered to be either *inside* if 𝐀 ⋅ 𝐁 < 0 or *outside* if 𝐀 ⋅ 𝐁 > 0. The condition |𝐀 ⋅ 𝐁| > 1 nm was used to account for the thickness of the vault shell and to filter out ions stuck at the vault surface. A crossing event was counted when an ion moved from the *outside* to the *inside* or vice versa between frame *n–2* and frame *n*, provided that the ion remained within the cylindrical region in both frames. Cumulative ion permeations were obtained by summing the number of inward and outward crossing events across the trajectory. The ion permeation data were recorded at each frame, with frames being recorded at 1.5 ns intervals.

## Structural analysis

Analysis of the inter-chains contacts was performed in ChimeraX^67^ (version 1.8) between residues 1-379 of 6 chains imposing a search of contact residues with at least 11 Å^2^ of buried area with a default probe radius of 1.4 Å. Atomic distances of alpha carbons between couples of residues Ala801, Gln678 and Asp39, between chains EB and YB as well as chains OB and UA were calculated in ChimeraX^67^ (1.8) and used to describe the diameter of the waist and cap-helix of the vault.

Final validation of the structure was performed in PHENIX^68^ (version 1.21). Model resolution and Model resolution range (Å) were calculated with the average value and range of values for the local resolution at FSC threshold 0.5 at atom positions using ChimeraX^67^ (version 1.8).

## Data availability

Cryo-EM maps are deposited in the Electron Microscopy Data Bank (EMDB) with the following accession codes EMD-53415, EMD-53423 and EMD-53440, for the vault in primed conformation, committed conformation and for the 39-mer half vault, respectively. Local refinement cryo-EM maps of the vault’s waist are deposited on EMDB with the following accession codes EMD-53438 and EMD-53439 for the vault in primed and committed conformation, respectively.

Atomic structures are deposited in the Protein Data Bank (PDB) with the following accession codes 9QW9 and 9QWQ for the human vault in primed and in committed conformation, respectively.

SAXS results are deposited in the Small Angle Scattering Biological Data Bank (SASDB) with accession code SASDXJ3.

## Author contributions

F.L. supervised the project and carried out the experiments; S.M. N.M. S.C. D.M. B.G. C.E. D.C. and G.T. contributed to sample preparation and data collection. K.P.R. S.C.L. planned and carried out the simulations with help from C.P.. F.L. K.P.R. S.C.L. and J.A. performed data analysis. F.L. P.T. A.D.

A.d.M. A.C. A.M. G.H. acquired funding and supervised the project. F.L. J.A. K.P.R. S.C.L. wrote the initial manuscript. All authors discussed the results and contributed to the final manuscript.

## ACKNOWLEDGEMENTS

We acknowledge the help of Gianni Frascotti and Camilla Pantaleoni from the University of Milano Bicocca for insights on their experimental work. Additionally, we would like to thank Matteo De March and Mattia Fanetti, from the University of Nova Gorica for their valuable advice. We are thankful to Roman Jerala and Jaka Snoj at the National Institute of Chemistry in Ljubljana for granting us access and setting up their SEC-MALS system for our samples. Additionally, we are grateful to Matic Kisovec at the National Institute of Chemistry in Ljubljana for his continuous help on maintaining cryoSPARC on the national HPC. We thank the Max Planck Computing and Data Facility for providing the computing resources to run the MD simulations.

## REFERENCES

1. Kedersha, N. L. & Rome, L. H. Preparative agarose gel electrophoresis for the purification of small organelles and particles. Analytical Biochemistry 156, 161–170 (1986).

2. Stephen, A. G. et al. Assembly of Vault-like Particles in Insect Cells Expressing only the Major Vault Protein. Journal of Biological Chemistry 276, 23217–23220 (2001).

3. Tanaka, H. & Tsukihara, T. Structural studies of large nucleoprotein particles, vaults. Proceedings of the Japan Academy Series B: Physical and Biological Sciences 88, 416–433 (2012).

4. Kato, K. et al. A vault ribonucleoprotein particle exhibiting 39-fold dihedral symmetry. Acta Crystallographica Section D: Biological Crystallography 64, 525 (2008).

5. Kedersha, N. L., Heuser, J. E., Chugani, D. C. & Rome, L. H. Vaults. III. Vault ribonucleoprotein particles open into flower-like structures with octagonal symmetry. Journal of Cell Biology 112, 225–235 (1991).

6. Tanaka, H. et al. The Structure of Rat Liver Vault at 3.5 Angstrom Resolution. Science 323, 384– 388 (2009).

7. Ma, C. et al. Structural insights into the membrane microdomain organization by SPFH family proteins. Cell Res 32, 176–189 (2022).

8. Fu, Z. & MacKinnon, R. Structure of the flotillin complex in a native membrane environment. Proc Natl Acad Sci U S A 121, e2409334121 (2024).

9. Ding, K. et al. Solution Structures of Engineered Vault Particles. Structure 26, 619–626.e3 (2018).

10. Casañas, A. et al. New features of vault architecture and dynamics revealed by novel refinement using the deformable elastic network approach. Acta Cryst D 69, 1054–1061 (2013).

11. Daly, T. K., Sutherland-Smith, A. J. & Penny, D. In silico resurrection of the major vault protein suggests it is ancestral in modern eukaryotes. Genome Biol Evol 5, 1567–1583 (2013).

12. Kedersha, N. L., Miquel, M. C., Bittner, D. & Rome, L. H. Vaults. II. Ribonucleoprotein structures are highly conserved among higher and lower eukaryotes. Journal of Cell Biology 110, 895–901 (1990).

13. Chugani, D. C., Rome, L. H. & Kedersha, N. L. Evidence that vault ribonucleoprotein particles localize to the nuclear pore complex. Journal of Cell Science 106, (1993).

14. Vollmar, F. et al. Assembly of nuclear pore complexes mediated by major vault protein. Journal of Cell Science 122, 780–786 (2009).

15. Chung, J. H., Ginn-Pease, M. E. & Eng, C. Phosphatase and tensin homologue deleted on chromosome 10 (PTEN) has nuclear localization signal-like sequences for nuclear import mediated by major vault protein. Cancer Research 65, 4108–4116 (2005).

16. Kim, E. et al. Crosstalk between Src and major vault protein in epidermal growth factor-dependent cell signalling. FEBS Journal 273, 793–804 (2006).

17. Kowalski, M. P. et al. Host resistance to lung infection mediated by major vault protein in epithelial cells. Science 317, 130–132 (2007).

18. Wang, W., Xiong, L., Wang, P., Wang, F. & Ma, Q. Major vault protein plays important roles in viral infection. IUBMB Life 72, 624–631 (2020).

19. Shimamoto, Y. et al. Direct activation of the human major vault protein gene by DNA-damaging agents. Oncology Reports 15, 645–652 (2006).

20. Ryu, S. J. et al. On the role of major vault protein in the resistance of senescent human diploid fibroblasts to apoptosis. Cell Death and Differentiation 15, 1673–1680 (2008).

21. Rayo, J. et al. Immunoediting role for major vault protein in apoptotic signaling induced by bacterial N-acyl homoserine lactones. Proc Natl Acad Sci U S A 118, e2012529118 (2021).

22. Berger, W., Steiner, E., Grusch, M., Elbling, L. & Micksche, M. Vaults and the major vault protein: Novel roles in signal pathway regulation and immunity. Cellular and Molecular Life Sciences 66, 43–61 (2009).

23. Frascotti, G. et al. The Vault Nanoparticle: A Gigantic Ribonucleoprotein Assembly Involved in Diverse Physiological and Pathological Phenomena and an Ideal Nanovector for Drug Delivery and Therapy. Cancers 13, 707 (2021).

24. Kickhoefer, V. A. et al. Vaults are up-regulated in multidrug-resistant cancer cell lines. Journal of Biological Chemistry 273, 8971–8974 (1998).

25. Izquierdo, M. A. et al. Broad distribution of the multidrug resistance-related vault lung resistance protein in normal human tissues and tumors. American Journal of Pathology 148, 877–887 (1996).

26. Huffman, K. E. & Corey, D. R. Major vault protein does not play a role in chemoresistance or drug localization in a non-small cell lung cancer cell line. Biochemistry 44, 2253–2261 (2005).

27. Mossink, M. H., Van Zon, A., Scheper, R. J., Sonneveld, P. & Wiemer, E. A. C. Vaults: A ribonucleoprotein particle involved in drug resistance? Oncogene 22, 7458–7467 (2003).

28. Kretz, P. F. et al. Dissecting the autism-associated 16p11.2 locus identifies multiple drivers in neuroanatomical phenotypes and unveils a male-specific role for the major vault protein. Genome Biology 24, 261 (2023).

29. Hananya, N., Ye, X., Koren, S. & Muir, T. W. A genetically encoded photoproximity labeling approach for mapping protein territories. Proceedings of the National Academy of Sciences 120, e2219339120 (2023).

30. Kedersha, N. L. & Rome, L. H. Isolation and characterization of a novel ribonucleoprotein particle: Large structures contain a single species of small RNA. Journal of Cell Biology 103, 699–709 (1986).

31. Kickhoefer, V. A. et al. The 193-kD vault protein, VPARP, is a novel poly(ADP-ribose) polymerase. Journal of Cell Biology 146, 917–928 (1999).

32. Cruz-León, S. et al. High-confidence 3D template matching for cryo-electron tomography. Nat Commun 15, 3992 (2024).

33. Poderycki, M. J. et al. The vault exterior shell is a dynamic structure that allows incorporation of vault-associated proteins into its interior. Biochemistry 45, 12184–12193 (2006).

34. Yang, J. et al. Vaults are dynamically unconstrained cytoplasmic nanoparticles capable of half vault exchange. ACS Nano 4, 7229–7240 (2010).

35. Llauró, A. et al. Decrease in pH destabilizes individual vault nanocages by weakening the inter-protein lateral interaction. Scientific Reports 6, 1–9 (2016).

36. Goldsmith, L. E., Yu, M., Rome, L. H. & Monbouquette, H. G. Vault Nanocapsule Dissociation into Halves Triggered at Low pH. ACS Publications 46, 2865–2875 (2007).

37. Querol-Audí, J. et al. The mechanism of vault opening from the high resolution structure of the N-terminal repeats of MVP. EMBO Journal 28, 3450–3457 (2009).

38. Guerra, P. et al. Symmetry disruption commits vault particles to disassembly. Science Advances 8, eabj7795 (2022).

39. Bernauer, L., Radkohl, A., Lehmayer, L. G. K. & Emmerstorfer-Augustin, A. Komagataella phaffii as Emerging Model Organism in Fundamental Research. Front. Microbiol. 11, (2021).

40. Galbiati, E. et al. A fast and straightforward procedure for vault nanoparticle purification and the characterization of its endocytic uptake. Biochimica et Biophysica Acta (BBA) - General Subjects 1862, 2254–2260 (2018).

41. Tomaino, G. et al. Addressing Critical Issues Related to Storage and Stability of the Vault Nanoparticle Expressed and Purified from Komagataella phaffi. Int J Mol Sci 24, 4214 (2023).

42. Punjani, A. & Fleet, D. J. 3D variability analysis: Resolving continuous flexibility and discrete heterogeneity from single particle cryo-EM. Journal of Structural Biology 213, 107702 (2021).

43. Punjani, A., Rubinstein, J. L., Fleet, D. J. & Brubaker, M. A. cryoSPARC: algorithms for rapid unsupervised cryo-EM structure determination. Nat Methods 14, 290–296 (2017).

44. Pei, J. & Grishin, N. V. AL2CO: calculation of positional conservation in a protein sequence alignment. Bioinformatics 17, 700–712 (2001).

45. Huiskonen, J. T. Image processing for cryogenic transmission electron microscopy of symmetry-mismatched complexes. Bioscience Reports 38, BSR20170203 (2018).

46. Trabuco, L. G., Villa, E., Mitra, K., Frank, J. & Schulten, K. Flexible fitting of atomic structures into electron microscopy maps using molecular dynamics. Structure 16, 673–683 (2008).

47. Croll, T. I. ISOLDE: a physically realistic environment for model building into low-resolution electron-density maps. Acta Cryst D 74, 519–530 (2018).

48. Kutzner, C. et al. Scaling of the GROMACS Molecular Dynamics Code to 65k CPU Cores on an HPC Cluster. Journal of Computational Chemistry 46, e70059 (2025).

49. Kimanius, D. & Schwab, J. Confronting heterogeneity in cryogenic electron microscopy data: Innovative strategies and future perspectives with data-driven methods. Current Opinion in Structural Biology 86, 102815 (2024).

50. Woodward, C. L., Mendonça, L. M. & Jensen, G. J. Direct visualization of vaults within intact cells by electron cryo-tomography. Cellular and Molecular Life Sciences: CMLS 72, 3401 (2015).

51. Yousefi, A. et al. Structural Flexibility and Disassembly Kinetics of Single Ferritin Molecules Using Optical Nanotweezers. ACS Nano 18, 15617–15626 (2024).

52. Sibarov, D. A. et al. Arc protein, a remnant of ancient retrovirus, forms virus-like particles, which are abundantly generated by neurons during epileptic seizures, and affects epileptic susceptibility in rodent models. Front Neurol 14, 1201104 (2023).

53. Teng, Y. et al. MVP-mediated exosomal sorting of miR-193a promotes colon cancer progression. Nat Commun 8, 14448 (2017).

54. Slinning, M. S. et al. Major vault protein is part of an extracellular cement material in the Atlantic salmon louse (Lepeophtheirus salmonis). Sci Rep 14, 15240 (2024).

55. Kickhoefer, V. A. et al. Engineering of vault nanocapsules with enzymatic and fluorescent properties. Proc Natl Acad Sci U S A 102, 4348–4352 (2005).

56. Doll, T. A. P. F., Raman, S., Dey, R. & Burkhard, P. Nanoscale assemblies and their biomedical applications. Journal of The Royal Society Interface 10, 20120740–20120740 (2013).

57. Muñoz-Juan, A., Carreño, A., Mendoza, R. & Corchero, J. L. Latest Advances in the Development of Eukaryotic Vaults as Targeted Drug Delivery Systems. Pharmaceutics 11, 300 (2019).

58. Tomaino, G. et al. An Efficient Method for Vault Nanoparticle Conjugation with Finely Adjustable Amounts of Antibodies and Small Molecules. Int J Mol Sci 25, 6629 (2024).

59. Wilkins, M. R. et al. Protein identification and analysis tools in the ExPASy server. *Methods in molecular biology (Clifton*, N.J.) 112, 531–52 (1999).

60. Franke, D. et al. ATSAS 2.8: A comprehensive data analysis suite for small-angle scattering from macromolecular solutions. Journal of Applied Crystallography 50, 1212–1225 (2017).

61. Grudinin, S., Garkavenko, M. & Kazennov, A. Pepsi-SAXS: an adaptive method for rapid and accurate computation of small-angle X-ray scattering profiles. *Acta crystallographica. Section D*, Structural biology 73, 449–464 (2017).

62. Marechal, N., Serrano, B. P., Zhang, X. & Weitz, C. J. Formation of thyroid hormone revealed by a cryo-EM structure of native bovine thyroglobulin. Nat Commun 13, 2380 (2022).

63. Vayssières, M. et al. Structural basis of DNA crossover capture by Escherichia coli DNA gyrase. Science 384, 227–232 (2024).

64. Scheres, S. H. W. RELION: implementation of a Bayesian approach to cryo-EM structure determination. J Struct Biol 180, 519–530 (2012).

65. Eswar, N. et al. Comparative Protein Structure Modeling Using MODELLER. Current Protocols in Protein Science 50, 2.9.1–2.9.31 (2007).

66. Emsley, P., Lohkamp, B., Scott, W. G. & Cowtan, K. Features and development of Coot. Acta Crystallogr D Biol Crystallogr 66, 486–501 (2010).

67. Meng, E. C. et al. UCSF ChimeraX: Tools for structure building and analysis. Protein Science 32, e4792 (2023).

68. Liebschner, D. et al. Macromolecular structure determination using X-rays, neutrons and electrons: recent developments in Phenix. Acta Crystallogr D Struct Biol 75, 861–877 (2019).

69. Afonine, P. V. et al. Real-space refinement in PHENIX for cryo-EM and crystallography. Acta Crystallogr D Struct Biol 74, 531–544 (2018).

70. Abraham, M. J. et al. GROMACS: High performance molecular simulations through multi-level parallelism from laptops to supercomputers. SoftwareX 1–2, 19–25 (2015).

71. Lindorff-Larsen, K. et al. Improved side-chain torsion potentials for the Amber ff99SB protein force field. Proteins 78, 1950–1958 (2010).

72. Berendsen, H. J. C., Postma, J. P. M., van Gunsteren, W. F., DiNola, A. & Haak, J. R. Molecular dynamics with coupling to an external bath. The Journal of Chemical Physics 81, 3684–3690 (1984).

73. Bussi, G., Donadio, D. & Parrinello, M. Canonical sampling through velocity rescaling. J Chem Phys 126, 014101 (2007).

74. Bernetti, M. & Bussi, G. Pressure control using stochastic cell rescaling. J Chem Phys 153, 114107 (2020).

75. de Jong, D. H. et al. Improved Parameters for the Martini Coarse-Grained Protein Force Field. J Chem Theory Comput 9, 687–697 (2013).

76. Kabsch, W. & Sander, C. Dictionary of protein secondary structure: pattern recognition of hydrogen-bonded and geometrical features. Biopolymers 22, 2577–2637 (1983).

77. Marrink, S. J., Risselada, H. J., Yefimov, S., Tieleman, D. P. & de Vries, A. H. The MARTINI force field: coarse grained model for biomolecular simulations. J Phys Chem B 111, 7812–7824 (2007).

78. Periole, X., Cavalli, M., Marrink, S.-J. & Ceruso, M. A. Combining an Elastic Network With a Coarse-Grained Molecular Force Field: Structure, Dynamics, and Intermolecular Recognition. J Chem Theory Comput 5, 2531–2543 (2009).

79. Javanainen, M., Martinez-Seara, H. & Vattulainen, I. Excessive aggregation of membrane proteins in the Martini model. PLoS One 12, e0187936 (2017).

80. Benayad, Z., von Bülow, S., Stelzl, L. S. & Hummer, G. Simulation of FUS Protein Condensates with an Adapted Coarse-Grained Model. J Chem Theory Comput 17, 525–537 (2021).

81. Humphrey, W., Dalke, A. & Schulten, K. VMD: visual molecular dynamics. J Mol Graph 14, 33– 38, 27–28 (1996).

82. Michaud-Agrawal, N., Denning, E. J., Woolf, T. B. & Beckstein, O. MDAnalysis: a toolkit for the analysis of molecular dynamics simulations. J Comput Chem 32, 2319–2327 (2011).

