## Supplementary Information for "Structural flexibility of the human vault protein revealed by high-resolution cryo-EM and molecular dynamics simulations"

[1] Laboratory for Environmental and Life Sciences, University of Nova Gorica, Vipavska cesta 13, 5000 Nova Gorica, Slovenia
[2] International Centre for Genetic Engineering and Biotechnology (ICGEB), Padriciano 99, 34149, Trieste, Italy
[3] Department of Theoretical Biophysics, Max Planck Institute of Biophysics, Max-von-Laue-Straße 3, 60438 Frankfurt am Main, Germany
[4] Consiglio Nazionale delle Ricerche - Istituto Officina dei Materiali c/o International School for Advanced Studies, via Bonomea 265, 34136 Trieste, Italy
[5] Institut de Génétique et de Biologie Moléculaire et Cellulaire (IGBMC), Parc D'Innovation 1 Rue Laurent Fries, 67404 Illkirch Cedex, France
[6] BM29 BIOSAXS beamline, European Synchrotron Radiation Facility (ESRF), Grenoble, France
[7] Istituto di Cristallografia, Consiglio Nazionale delle Ricerche, Strada Statale 14 km 163.5, 34149 Trieste, Italy
[8] Max Planck Tandem Group Biophysics of Tropical Diseases, Faculty of Exact and Natural Sciences, University of Antioquia, 050010 Medellín, Colombia
[9] Department of Biotechnology and Biosciences, University of Milano-Bicocca, 20126 Milano, Italy
[10] Current address: IEO, European Institute of Oncology IRCCS, Department of Experimental Oncology, 20139 Milan, Italy
[11] Institute of Biophysics, Goethe University Frankfurt, 60438 Frankfurt am Main, Germany

### Table of Contents

Supplementary Figures 1 to 14
Supplementary Tables 1 and 2
Description of Supplementary Movies 1 and 2

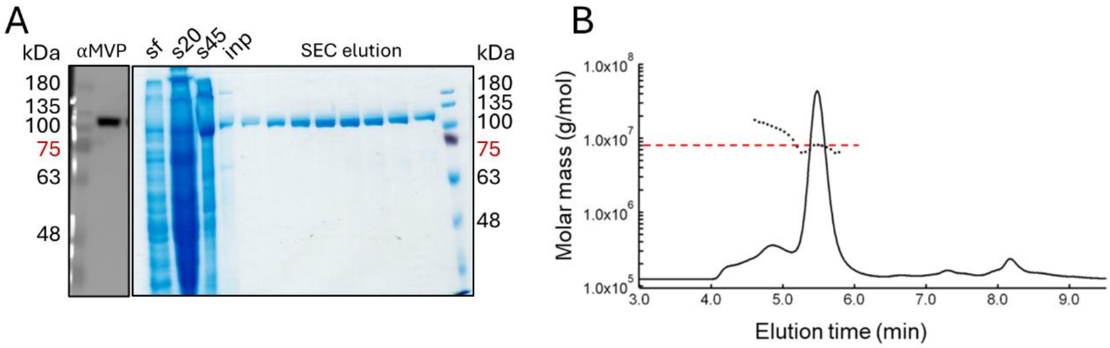


**Supplementary Figure 1 – Protein characterization.** A) Left: western blot of the protein sample, developed using anti-MVP antibody. Right: SDS-PAGE of the sample. Lanes from left to right: soluble fraction of the lysate (sf); supernatant after 20,000 x *g* centrifugation (s20); sucrose layer 45% (s45); SEC input after RNase incubation (inp); SEC fractions containing fully assembled vault. B) SEC coupled with Multi Angle Light scattering (SEC-MALS) analysis of the sample. The main peak corresponds to 7.66 ± 0.6% MDa. The UV signal at 280 nm is shown as black line and the molecular mass, calculated with 8 angular detectors, is represented by scattered points near the main peak (red dashed line at 8.0 MDa).

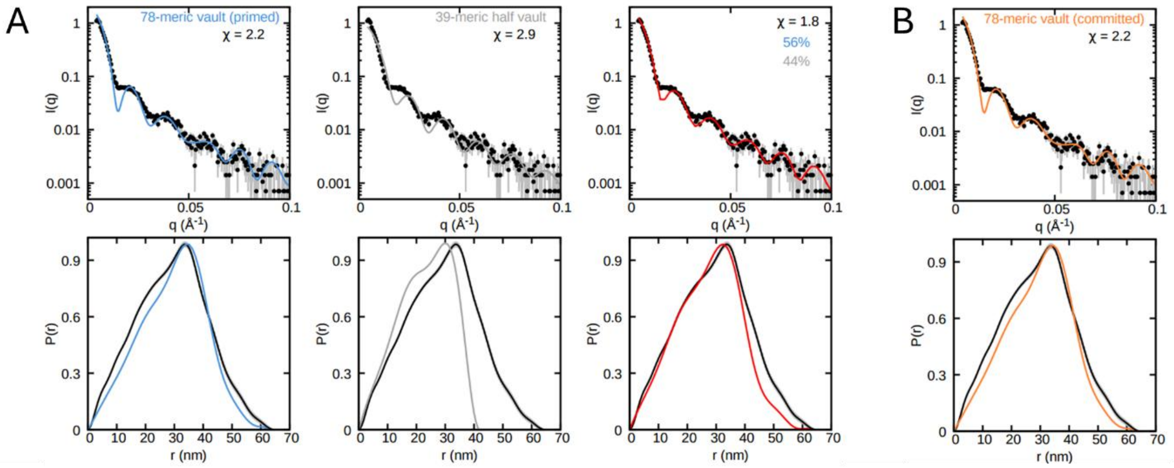


**Supplementary Figure 2 – SAXS sample analysis**. A) Experimental SAXS scattering curve and derived distance-distribution plot in black with fits to molecular models. The $\chi$ value represents the fit quality for: the 78-mer vault in primed conformation (in blue), the 39-mer half vault (in grey) or the calculated ideal ratio of 56% and 44% for the combination of 78-mer vault and the 39-mer half-vault, respectively (in red). B) Experimental SAXS scattering curve and derived distance-distribution plot in black with fits to the molecular model of the 78-mer vault in the committed conformation (in orange).

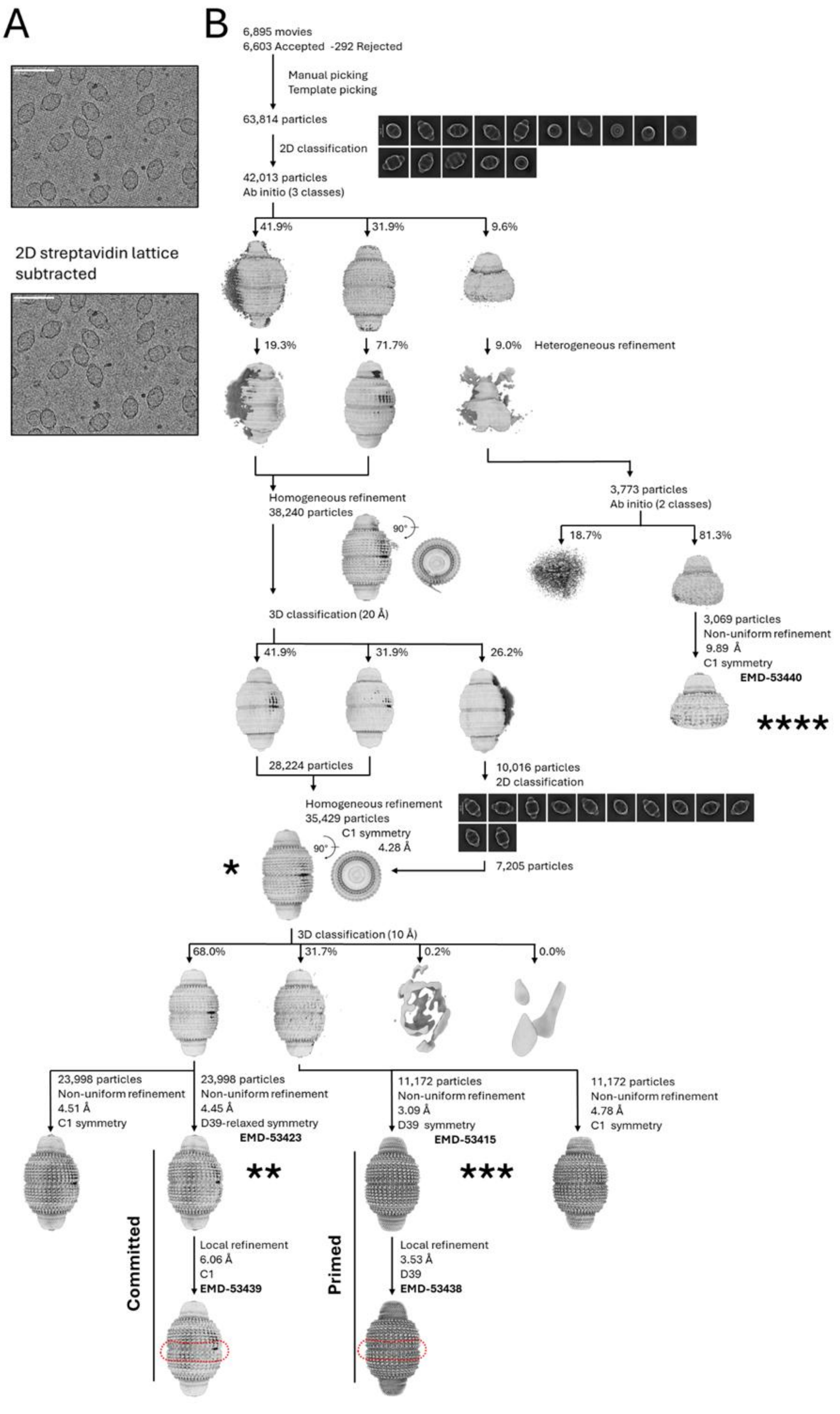
A
B
2D streptavidin lattice subtracted
6,895 movies
6,603 Accepted -292 Rejected
Manual picking
Template picking
63,814 particles
2D classification
42,013 particles
Ab initio (3 classes)
41.9%
31.9%
9.6%
19.3%
71.7%
9.0%
Heterogeneous refinement
Homogeneous refinement
38,240 particles
90°
3,773 particles
Ab initio (2 classes)
18.7%
81.3%
3,069 particles
Non-uniform refinement
9.89 Å
C1 symmetry
EMD-53440
****
3D classification (20 Å)
41.9%
31.9%
26.2%
28,224 particles
10,016 particles
2D classification
Homogeneous refinement
35,429 particles
C1 symmetry
4.28 Å
90°
*
7,205 particles
3D classification (10 Å)
68.0%
31.7%
0.2%
0.0%
23,998 particles
Non-uniform refinement
4.51 Å
C1 symmetry
23,998 particles
Non-uniform refinement
4.45 Å
D39-relaxed symmetry
EMD-53423
11,172 particles
Non-uniform refinement
3.09 Å
D39 symmetry
EMD-53415
11,172 particles
Non-uniform refinement
4.78 Å
C1 symmetry
**
***
Committed
Local refinement
6.06 Å
C1
EMD-53439
Primed
Local refinement
3.53 Å
D39
EMD-53438

**Supplementary Figure 3 – cryo-EM data processing**. A) Above, representative cryo-EM micrograph. Below, micrograph after subtraction of the streptavidin 2D crystal lattice by Fourier filtering of Bragg spots. B) Flowchart of data processing performed in cryoSPARC. The procedure used, particle numbers, enforced symmetry, final resolution and eventual region of the mask used for local refinement (in red dotted line) are indicated at each step in the flowchart. Unsharpened maps from the 3D refinement passages and representative 2D classes from reference-free 2D classification are reported. *) initial refinement of the vault, then classified into the two conformational states. **) Committed conformation of the vault obtained with relaxed D39 symmetry. ***) primed conformation of the vault obtained with D39 symmetry. ****) 39-mer half-vault.

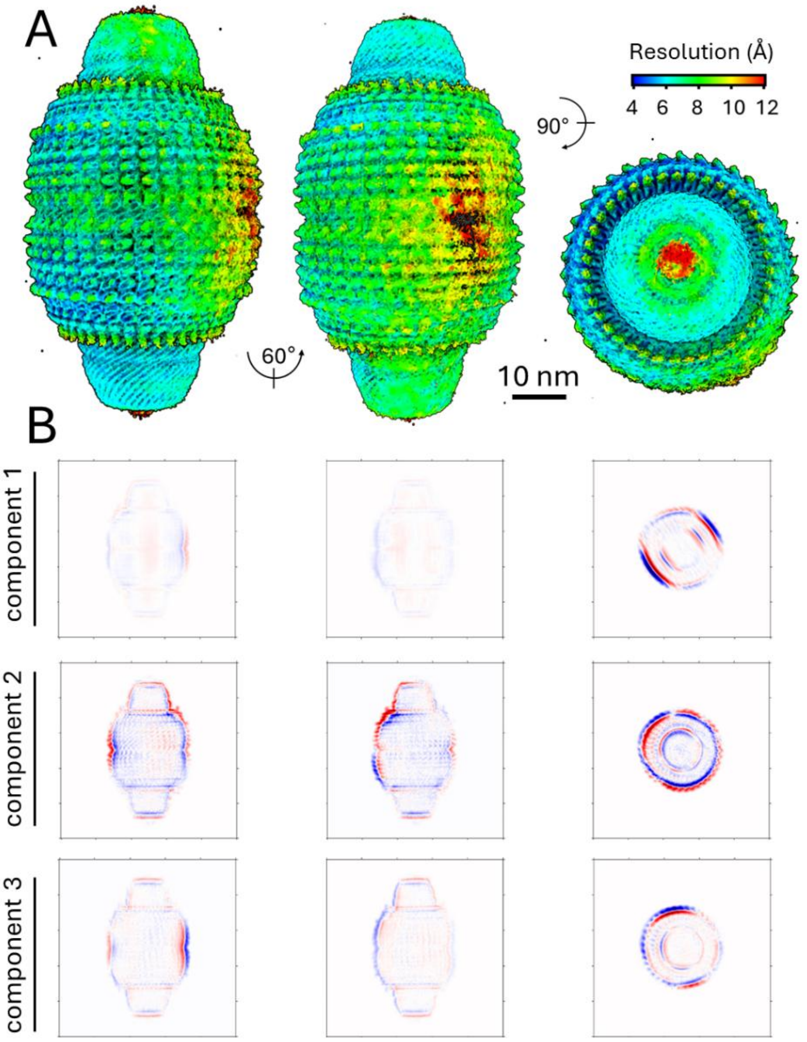


**Supplementary Figure 4 – Initial reconstruction of the vault particle.** A) Volume resulting from homogeneous refinement of the 35,429 particles identified as intact vaults. The local resolution of the map is shown in rainbow color scale. B) Three orthogonal slices through each variability component of the 3D volume resulting from 3D variability analysis (3DVA) applied to the same 35,429 particles (see also Supplementary Movie 1). To explain heterogeneity within the particle stack, positive and negative values in the 3D volume at each voxel are shown in a color scale from blue to red, respectively.

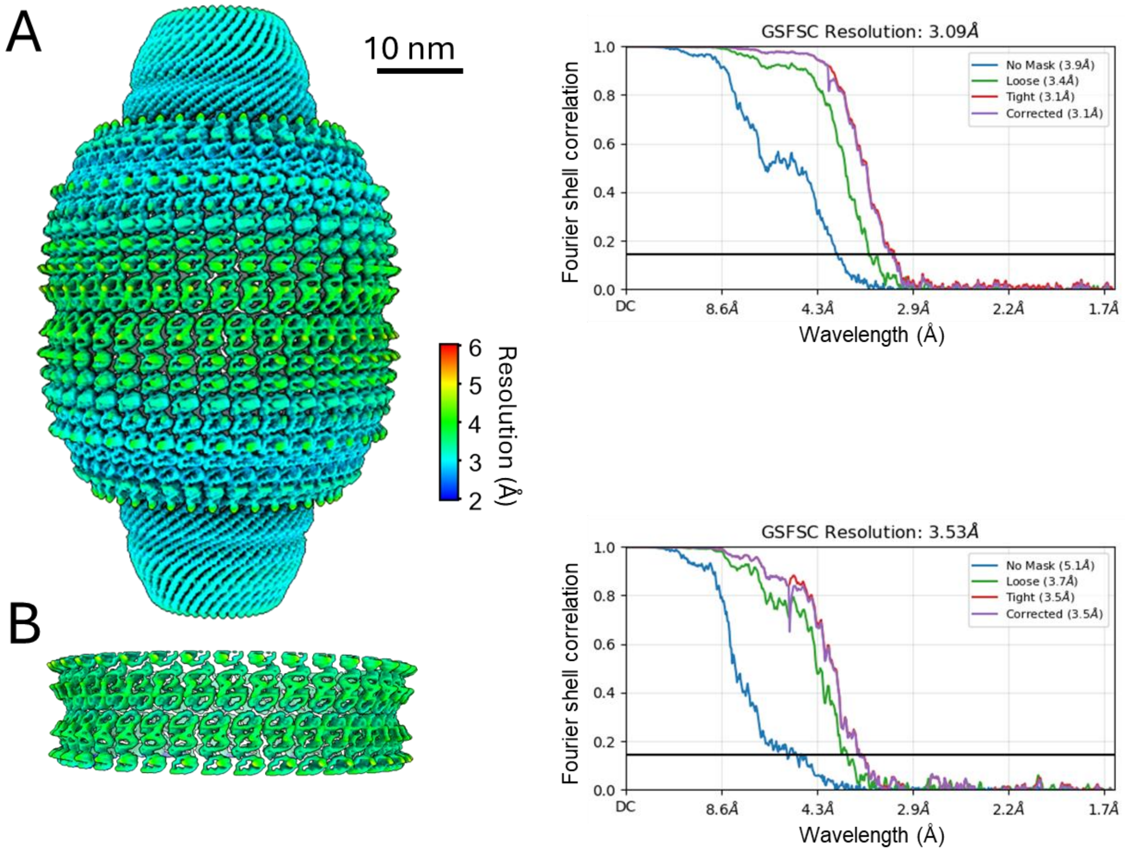


**Supplementary Figure 5 – Reconstructed maps of the vault in primed conformation.** A) Local resolution of the map (unsharpened) resulting from the cryo-EM reconstruction of the vault in the primed conformation, shown in rainbow color scale. The corresponding GS-FSC curve is displayed on the right. B) Local resolution of the map (unsharpened) resulting from the cryo-EM local reconstruction of the vault's waist in the primed conformation, also shown in rainbow color scale. The corresponding GS-FSC curve is displayed on the right.

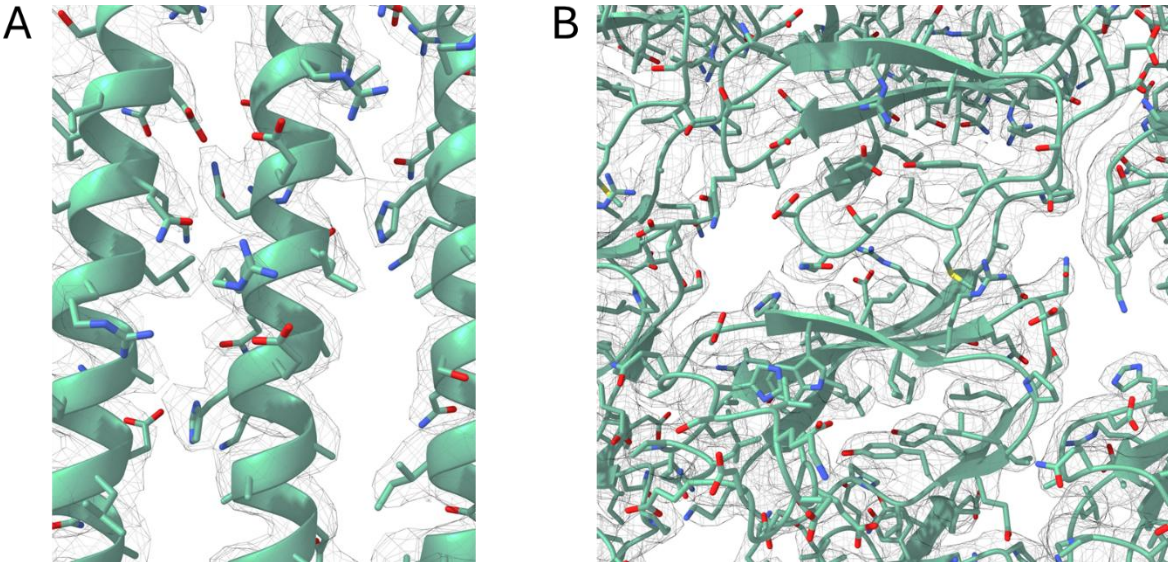


**Supplementary Figure 6 – Side chain resolution of the primed vault**. A) Detail of sharpened map of the cap-helix of the vault in the primed conformation (contour level 0.8). B) Detail of the sharpened map of the repeat domains of the vault in the primed conformation (contour level 0.8).

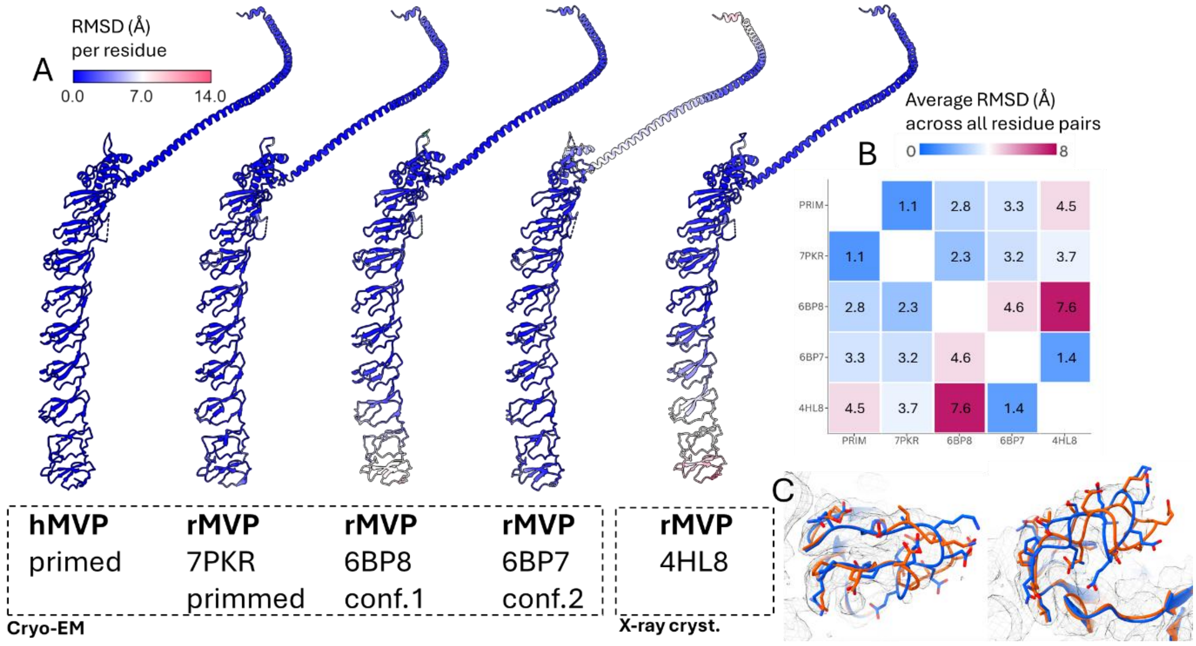


**Supplementary Figure 7 – Structural comparison of the human vault to murine structures**. A) RMSD between atomic coordinates of the primed human vault (hMVP) and previously published *Rattus norvegicus* vault (rMVP) structures. B) RMSD between all atom pairs in hMVP and all published rMVP structures. C) Detailed structural comparison of the loop 342-347, showing the hMVP primed conformation (this work) in blue, overlaid with the crystal structure (PDB: 4HL8) in red. The meshed cryo-EM density map (contour level 0.8) from this work is also shown.

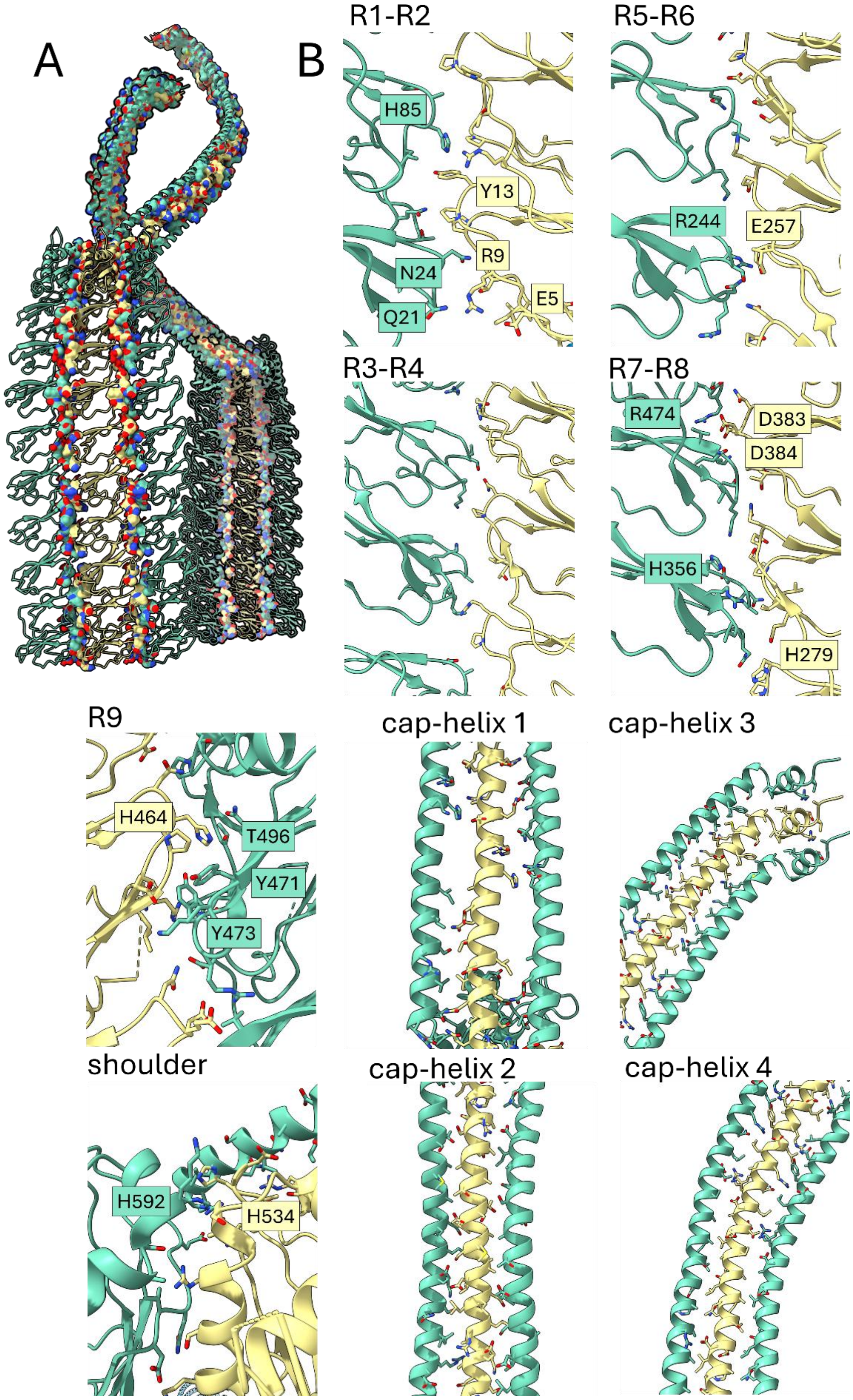


**Supplementary Figure 8 – Neighboring lateral contacts between MVP monomers in the primed vault.** A) Cartoon representation of three MVP monomers, with surface view of residues involved in neighboring contacts between laterally interacting subunits. B) Detailed views of different regions of the atomic model of the vault in the primed conformation, represented as cartoon. Side chains of residues involved in lateral contacts are shown as sticks. Ionizable residues involved in polar interactions are labelled.

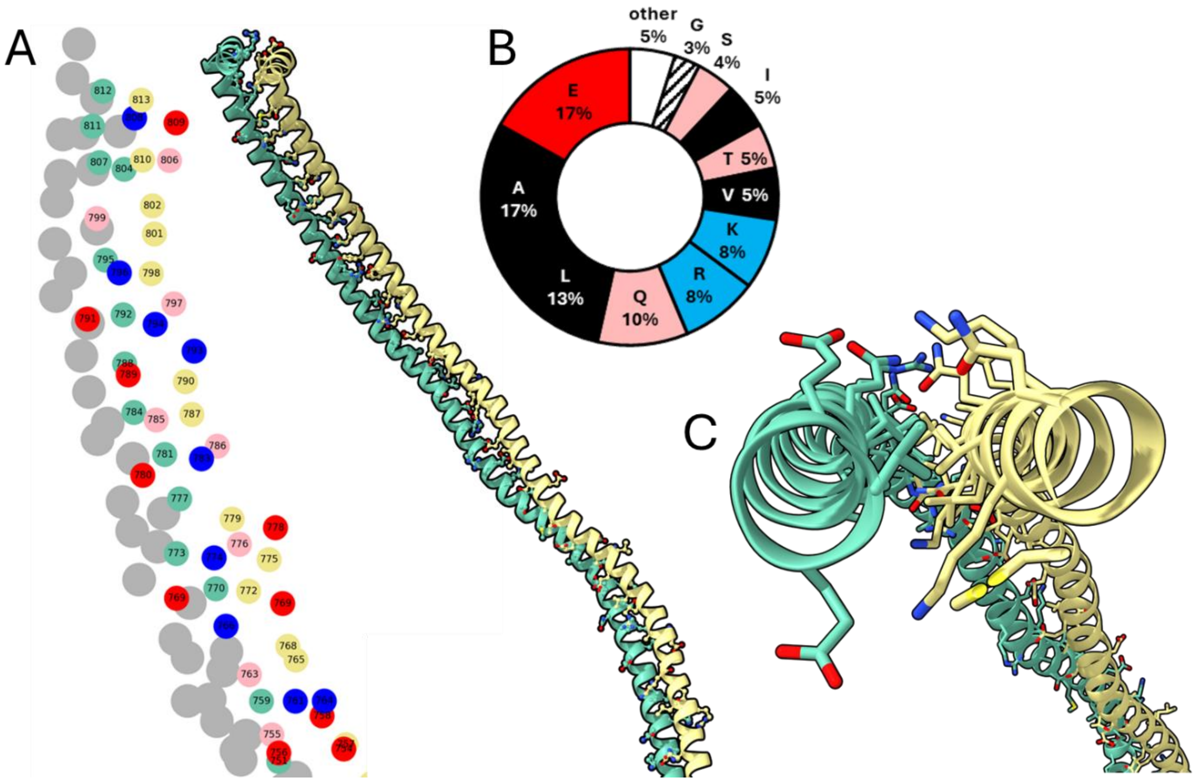


**Supplementary Figure 9 – Lateral contacts in the helix-cap domain of hMVP.** A) The residues' number of amino acids involved in lateral contacts between two MVP chains in circles, with blue and red indicating positively and negatively charged residues, respectively, yellow and green for other residues. The plot represents a side view of the atomic model, shown on the right. B) Pie chart, showing the distribution of residues found in later chains MVP-MVP contacts. C) Top view of two helices in the cap-helix, showing knobs-into-holes contacts established by the inner cleft residues between the two helices and the polar residues surrounding the hydrophobic cleft.

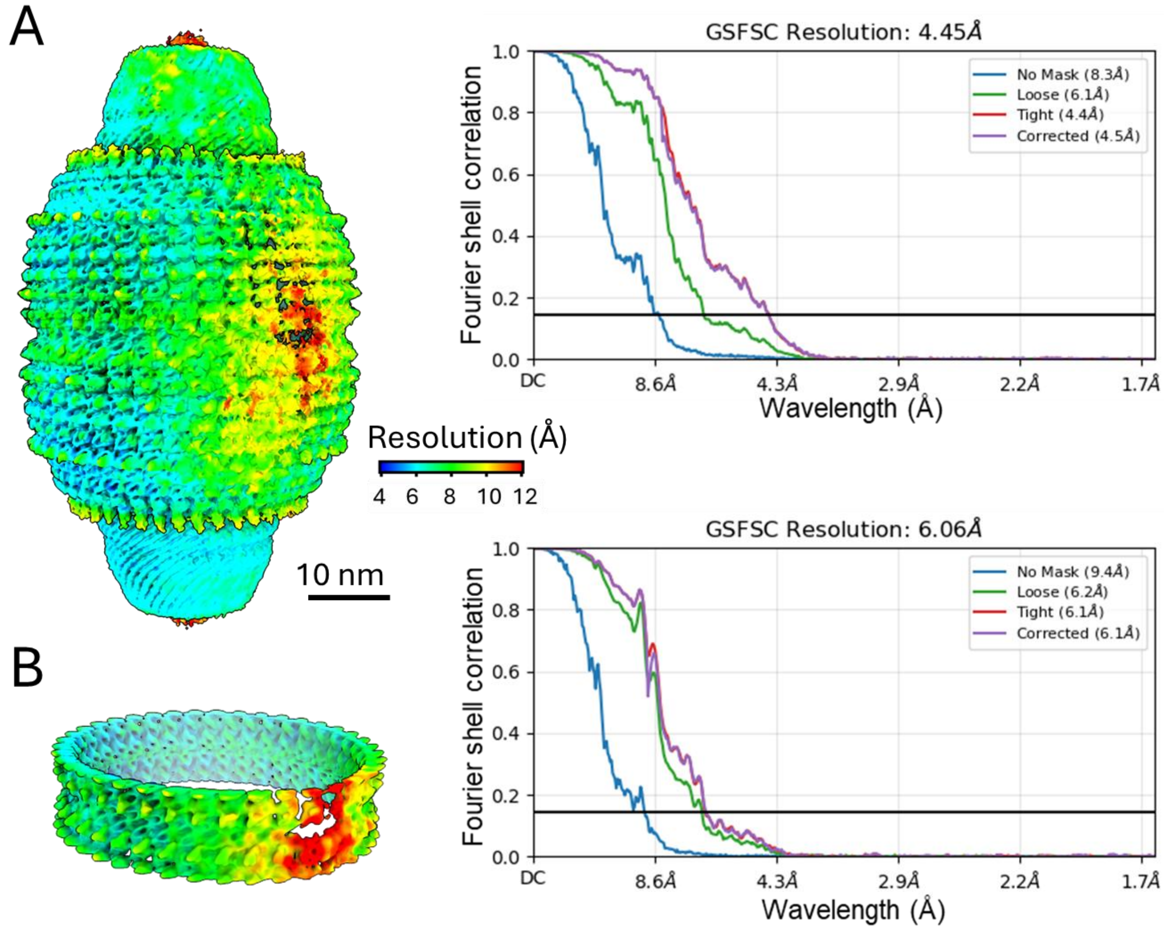


**Supplementary Figure 10 – Refined map of the vault in committed conformation**. A) Local resolution map of the vault in the committed conformation, obtained from non-uniform refinement of 23,998 particles. The corresponding GS-FSC (gold-standard Fourier shell correlation) curve is shown on the right, calculated between two independent half-maps using cryoSPARC, with the resolution values reported at 0.143 FSC. B) Resolution estimation of the local refinement of the vault's waist, with the corresponding GS-FSC curve displayed on the right, calculated between two independent half-maps using cryoSPARC, with the resolution values reported at 0.143 FSC.

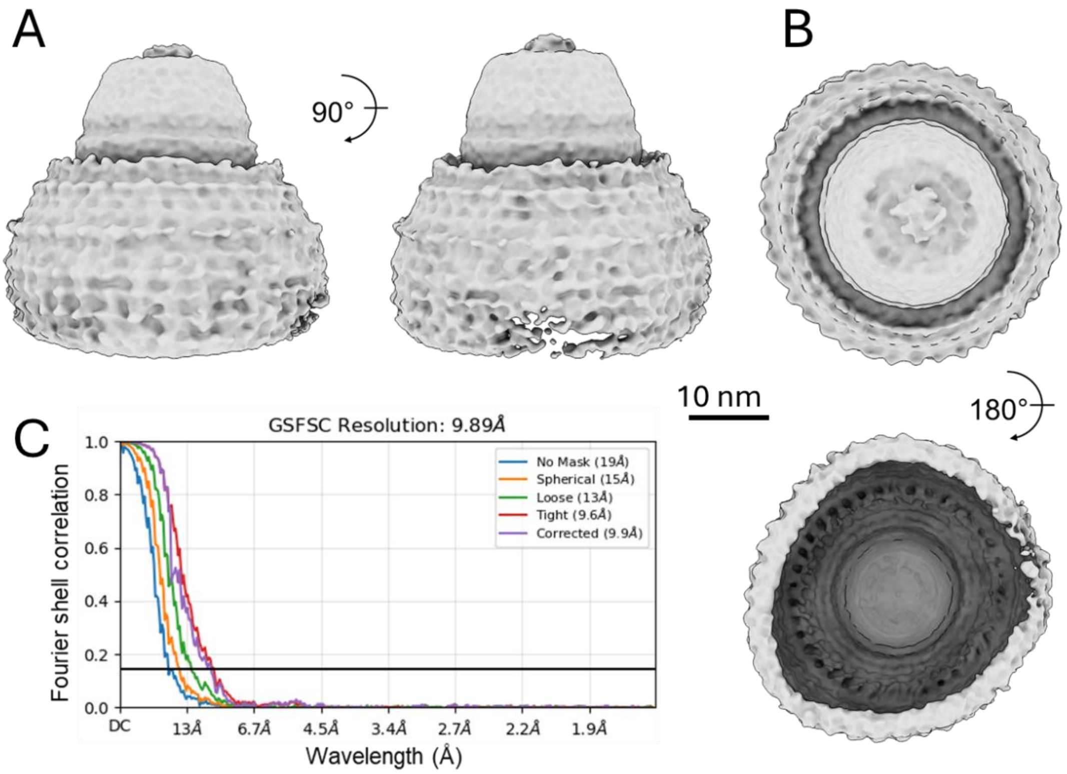


**Supplementary Figure 11 – Refinement of the 39-mer half vault**. A) Lateral views of the non-uniform refinement of the 39-mer half vault, obtained from 3,069 particles (contour level 0.3). B) Top and bottom views of the non-uniform refinement of the 39-mer half vault. C) Corresponding GS-FSC curve, calculated between two independent half-maps using cryoSPARC, with resolution values reported at 0.143 FSC.

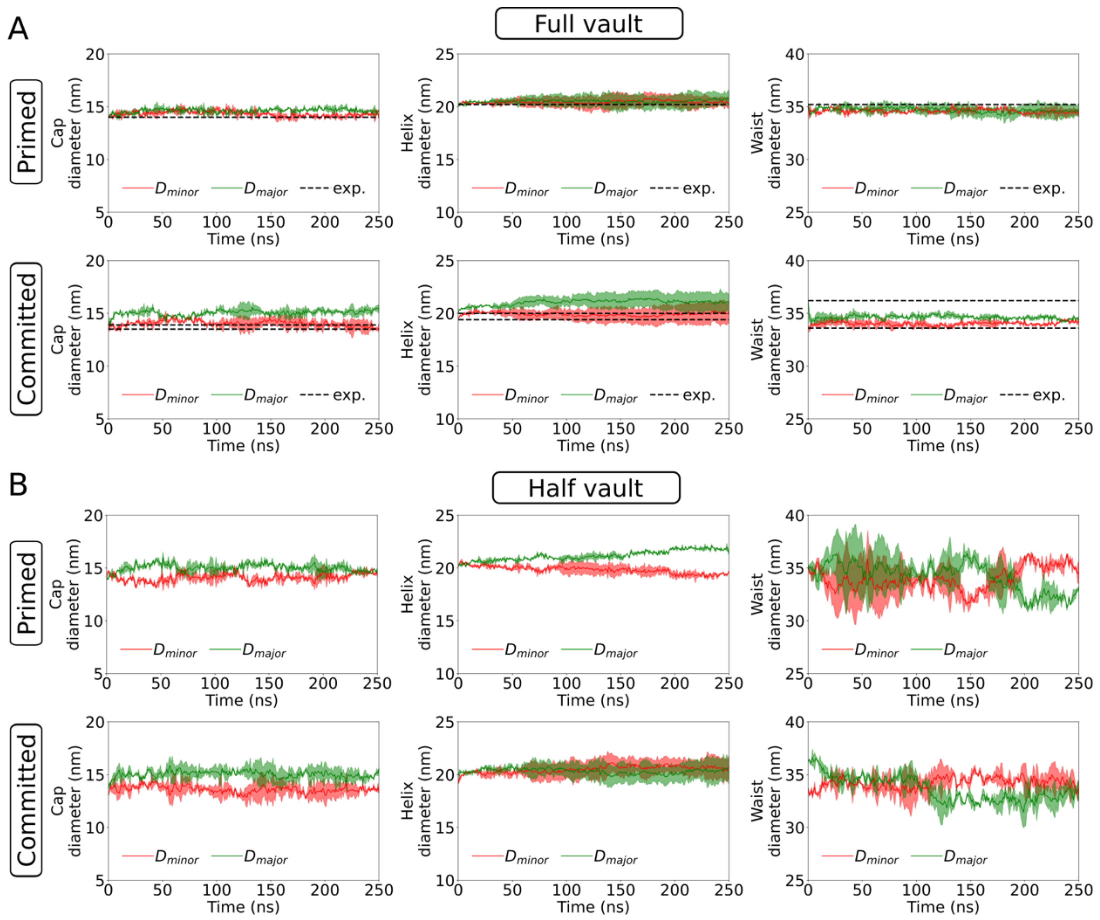


**Supplementary Figure 12 – Diameter of cap, helix and waist regions in AA-MD simulations across vault conformations.** A) Time series of the vault's cap (left), helix (center), and waist (right) diameters in the primed (top) and committed (bottom) conformations for the full vault. Diameters are obtained by fitting an ellipse to the backbone bead positions of residues A801 (cap), Q678 (helix), and D39 (waist), with major and minor diameters shown in green and red, respectively. Mean values (solid lines) and two standard deviations (shaded regions) from two simulation replicates are plotted. Dashed lines indicate diameters from the atomic model. B) Time series of the vault's cap (left) and helix (right) diameters in the primed (top) and committed (bottom) conformations for the half vault, estimated in the same way as in panel A.

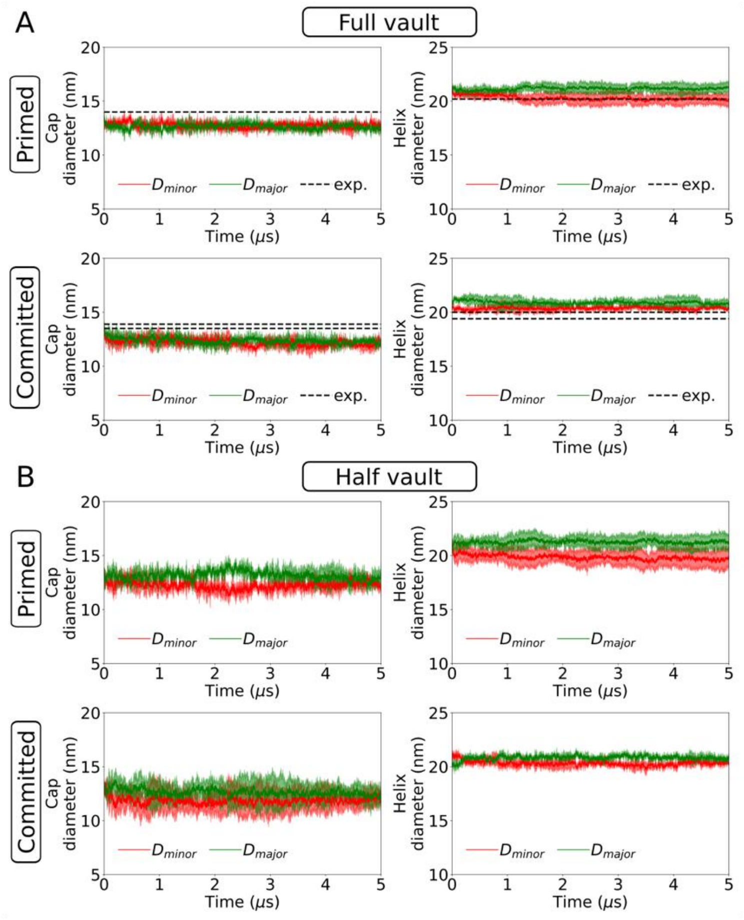


**Supplementary Figure 13 – Diameter of cap and helix regions in CG-MD simulations across vault conformations.** A) Time series of the vault's cap (left) and helix (right) diameters in the primed (top) and committed (bottom) conformations for the full vault. Diameters are obtained by fitting an ellipse to the backbone bead positions of residues A801 (cap) and Q678 (helix), with major and minor diameters shown in green and red, respectively. Mean values (solid lines) and two standard deviations (shaded regions) from three simulation replicates are plotted. Dashed lines indicate diameters from the atomic model. B) Time series of the vault's cap (left) and helix (right) diameters in the primed (top) and committed (bottom) conformations for the half vault, estimated in the same way as in panel A.

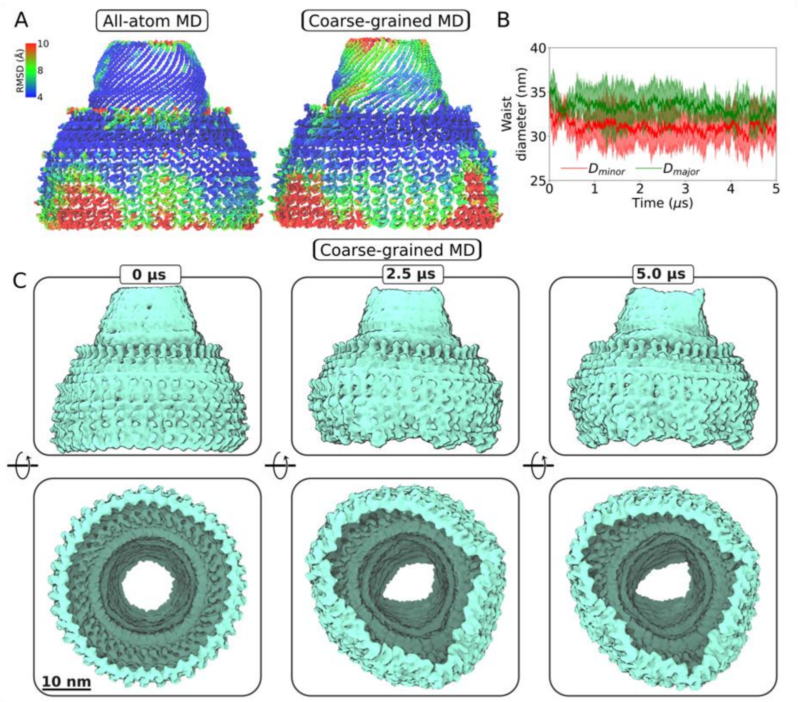


**Supplementary Figure 14 – All-atom (A) and coarse-grained (B–C) simulations of the half-vault in the primed conformation.** A) Left panel, RMSD over the last 200 ns of simulation, mapped onto the initial atomic structure of the half-vault in the primed conformation. Structures are aligned to the Cα atoms of residues with RMSF <3.5 Å. Right panel, RMSD over the last 4.5 μs of simulation, mapped onto the initial coarse-grained structure of the half-vault in the primed conformation. Structures are aligned to the backbone beads of residues with RMSF <3.5 Å. B) Time series of the half vault's waist diameter in the primed conformation. Diameters are obtained by fitting an ellipse to the backbone bead positions of residues D39, with major and minor diameters shown in green and red, respectively. Mean values (solid lines) and two standard deviations (shaded regions) from three simulation replicates are plotted. C) Coarse-grained MD simulations of the half-vault in the primed conformation. Representative snapshots from the MD trajectory are shown from a lateral view (top) and an internal bottom-up view (bottom). The coarse-grained structures are represented as surfaces (contour level 1.3).

**Supplementary Table 1 – Cryo-EM data collection, refinement and validation statistics**

| | Vault primed (EMD-53415) (PDB 9QW9) | Local refin. waist primed vault (EMD-53438) | Vault committed (EMD-53423) (PDB 9QWQ) | Local refin. waist committed vault (EMD-53439) | 39-mer half vault (EMD-53440) |
|---|---|---|---|---|---|
| **Data collect. and processing** | | | | | |
| Magnification | 81,000 | 81,000 | 81,000 | 81,000 | 81,000 |
| Voltage (kV) | 300 | 300 | 300 | 300 | 300 |
| Electr. expos. (e–/Å$^2$) | 50.15 | 50.15 | 50.15 | 50.15 | 50.15 |
| Defocus range (μm) | 0.55-2.33 | 0.55-2.33 | 0.55-2.33 | 0.55-2.33 | 0.55-2.33 |
| Pixel size (Å) | 0.8416 | 0.8416 | 0.8416 | 0.8416 | 0.8416 |
| Symmetry imposed | D39 | D39 | Relaxed D39 | C1 | C1 |
| Initial particles | 63,814 | 63,814 | 63,814 | 63,814 | 63,814 |
| Final particles | 11,172 | 11,172 | 23,998 | 23,998 | 3,069 |
| Map resolution (Å) | 3.09 | 3.53 | 4.45 | 6.06 | 9.89 |
| FSC threshold | 0.143 | 0.143 | 0.143 | 0.143 | 0.143 |
| Map resolution. range (Å) | 26.98 – 2.23 | 5.29 – 2.64 | 46.55 – 3.84 | 12.20 – 4.65 | 24.7 – 8.69 |
| **Refinement** | | | | | |
| Initial model used (PDB code) | 4HL8 | | 4HL8 | | |
| Model resolution (Å) | 3.1 | | 6.4 | | |
| FSC threshold | 0.5 | | 0.5 | | |
| Model resolution range (Å) | 5.67 – 2.68 | | 46.02 – 3.84 | | |
| Map sharpening *B* factor (Å$^2$) | -144.4 | | -110.9 | | |
| Model composition | | | | | |
| Non-hydrogen atoms | 482,118 | | 482,118 | | |
| Protein residues | 60,762 | | 60,762 | | |
| *B* factors (Å$^2$) | | | | | |
| Protein | 47.7/200.6/105.0 | | 169.7/1051.29/386.74 | | |
| R.m.s. deviations | | | | | |
| Bond lengths (Å) | 0.004 (0) | | 0.004 (1) | | |
| Bond angles (°) | 0.941 (0) | | 0.958 (11) | | |
| Validation | | | | | |
| MolProbity score | 1.92 | | 2.43 | | |
| Clashscore | 5.31 | | 18.29 | | |
| Poor rotamers (%) | 1.78 | | 0.00 | | |
| Ramachandran plot | | | | | |
| Favored (%) | 93.07 | | 84.50 | | |
| Allowed (%) | 6.85 | | 15.42 | | |
| Disallowed (%) | 0.08 | | 0.08 | | |

**Supplementary Table 2 – Description of the different MD production simulations reported in this work**

| System | Conformation | Type of MD | # replicates | Duration [ns] | Box size [nm] | # particles |
|---|---|---|---|---|---|---|
| Vault particle | Primed | AA-MD | 2 | 250 | 78.1x45.5x52.5 | 18358261 |
| Vault particle | Committed | AA-MD | 2 | 250 | 78.3x46.0x51.7 | 18363369 |
| Vault particle | Primed | CG-MD | 3 | 5000 | 77.9x45.3x52.3 | 1555766 |
| Vault particle | Committed | CG-MD | 3 | 5000 | 78.0x45.8x51.5 | 1560076 |
| Half vault particle | Primed | AA-MD | 2 | 250 | 52.5x45.5x45.2 | 10611232 |
| Half vault particle | Committed | AA-MD | 2 | 250 | 52.8x44.7x46.0 | 10710501 |
| Half vault particle | Primed | CG-MD | 3 | 5000 | 52.3x45.3x45.0 | 893571 |
| Half vault particle | Committed | CG-MD | 3 | 5000 | 52.5x44.5x45.7 | 896709 |

#### Description of Supplementary Movies

File Name: Supplementary Movie 1
Description: 3D Variability Analysis of the initial stack of 35,429 vault particles. The different trajectories (modes) in 3D volumes show where there is significant variability in the dataset.

File Name: Supplementary Movie 2
Representative trajectory of the CG-MD simulations. Vault in primed (left) and committed (right) conformations. The coarse-grained structures are shown as surfaces (contour level 1.3).
